## Supplementary Methods for "KSTAR: An algorithm to predict patient-specific kinase activities from phosphoproteomic data"

### Supplementary Methods: Expanded description of KSTAR algorithm

#### Content Found In Document


1. **Problems Addressed by the KSTAR Algorithm**
    (a) Issues with Kinase-Substrate Networks
        i. Problem 1: Kinase-substrate annotations are sparse (page 2)
        ii. Problem 2: Using thresholded kinase-substrate prediction networks degrades performance (page 2)
        iii. Problem 3: Thresholding kinase-substrate prediction networks results in "Hub" substrates (page 3)
        iv. Problem 4: Thresholding kinase-substrate prediction networks results in "Hub" kinases and kinases with little associated evidence (page 3-4)
        v. Problem 5: Certain kinases tend to be connected to well studied sites, leading to kinase-specific false positive rates. (page 3-4)
        vi. Problem 6: Kinases from the same family often exhibit high network overlap (connected to the same substrates) in thresholded networks (page 5)
    (b) Challenges with applying kinase activity inference
        i. Problem 7: Most kinase-activity inference algorithms rely on relative quantification (page 5)
        ii. Problem 8: Phosphoproteomic experiments tend to identify well-studied sites, leading to experiment specific false positive rates (page 6)
2. **Implementing the KSTAR algorithm**
    (a) Heuristic Prune and Generation of Network Ensemble
        i. Prune algorithm without accounting for kinase-specific study bias (page 6-7)
        ii. Prune algorithm when accounting for kinase-specific study bias (page 7-8)
    (b) Inferring Kinase Activity from Phosphoproteomic Data
        i. Calculating kinase-substrate enrichment in each pruned network (page 8-9)
        ii. Generating random phosphoproteomic experiments (page 9)
        iii. Calculating the final KSTAR activity score (page 9-10)
        iv. Calculating the false positive rate of the activity score (page 10)

#### Problems Addressed by the KSTAR Algorithm

##### Issues with Kinase-Substrate Networks

**Problem 1:** Known kinase-substrate annotations from phosphoproteome databases like PhosphoSitePlus (Hornbeck, Nucleic Acids Research, 2012) are sparse: only a small fraction of the total phosphoproteome have at least one known connection to a kinase.

**Solution:** Using information from kinase-substrate prediction networks like NetworKIN (Horn, 2014) would allow for better coverage of the phosphoproteome.

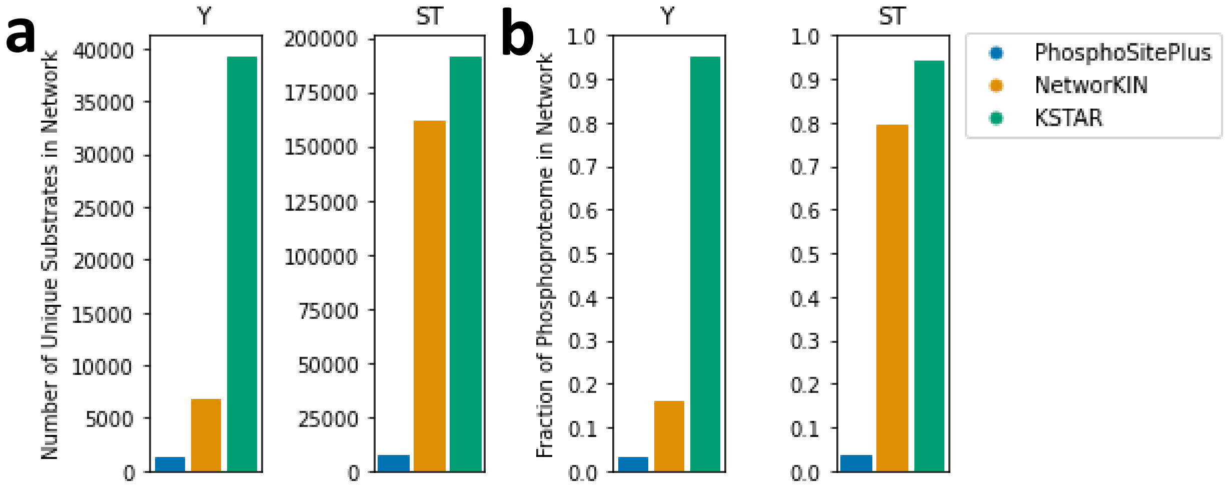


**Figure SM.1. KSTAR Expands the Number of Phosphorylation Sites Used in Prediction** A) The total number of unique substrates with a edge/interaction with at least one kinase for known kinase-substrate annotations in PhosphoSitePlus, or predicted interactions from a thresholded NetworKIN, or predicted interactions from KSTAR networks. B) The same information as A, but provided as a fraction of the total phosphoproteome.

**Problem 2:** Thresholded kinase-substrate prediction networks have been shown to exhibit poor performance relative to using kinase-substrate annotations alone.

**Solution:** We hypothesized there is useful information in these kinase-substrate networks, but that the single kinase-substrate network generated by thresholding contains too many incorrect predictions and issues with study bias to be useful. Instead, we proposed that we could create an ensemble of many possible kinase-substrate networks using the network edge weights to guide edge selection, rather than relying on single representation of kinase-substrate relationships (heuristic prune, Algorithm 2). This approach is built on the idea that all networks are wrong, but they are wrong in different ways. While any one single network is unlikely to be the correct representation, aggregating information across these networks will allow us to converge on the kinases most likely to be active. We have demonstrated the utility of ensemble approaches in some of our prior work, such as for clustering (OpenEnsembles, Ronan, Jornal of Machine Learning Research, 2018).

**Problem 3:** Thresholded kinase-substrate prediction networks result in the emergence of “Hub” substrates, which provide evidence for a disproportionate number of kinases.

**Solution:** During the heuristic prune, we limit the number of edges each substrate can have in each network (number of kinases it can provide evidence for).

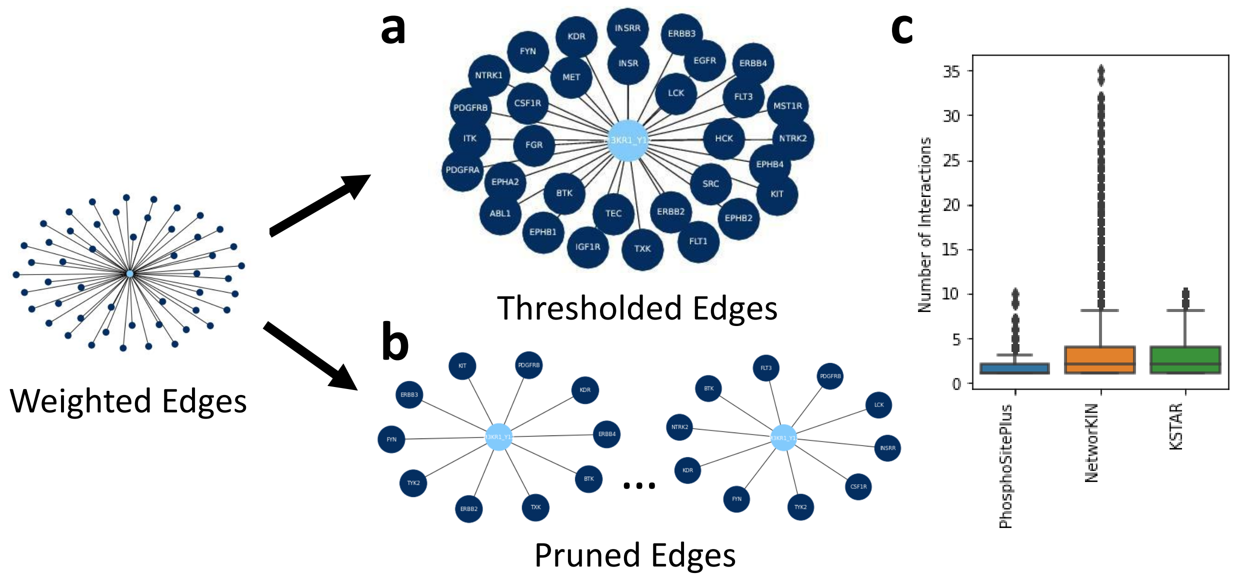


**Figure SM.2. Effect of Heuristic Prune on Hub Substrates in NetworKIN** Here, we demonstrate the impact of the prune on substrate hubs, using PI3KR1 Y12 as an example. A) Example hub substrate in NetworKIN thresholded with a value of 1 and B) the same substrate in two of the 50 pruned networks. C) Number of kinases each substrate is connected to in PhosphoSitePlus, NetworKIN thresholded with a value of 1, and KSTAR pruned networks. Thresholded NetworKIN and pruned KSTAR networks exhibit similar distributions, where the median number of kinases each substrate is connected to is 2. However, KSTAR networks do not have any outlier substrates connected to more than 10 kinases (“Hub” substrates), which can have a large influence on the final predictions.

**Problem 4:** Thresholded kinase-substrate prediction networks result in the emergence of “Hub” kinases, which are connected to a disproportionate number of substrates. In addition, many kinases often lose the majority of their edges in the network, losing the ability to generate predictions for these kinases.

**Solution:** During the heuristic prune, force all kinases to have the same number of edges (connections to substrates) in each network, ensuring that no one kinase has too few or too many substrates providing evidence for them.

**Problem 5:** Kinase-substrate prediction networks exhibit high study bias, where certain kinases are more likely to be connected to well-studied sites, which leads to kinase-specific false positive rates.

**Solution:** Enforce a rule during the heuristic prune that ensures every kinase has edges with the same distribution of study bias, as defined by the number of phosphoproteome compendia they are identified in. This means that no one kinase is connected to more well studied sites (site found in most compendia) and no one kinase is connected to more poorly studied sites (site found in few compendia).

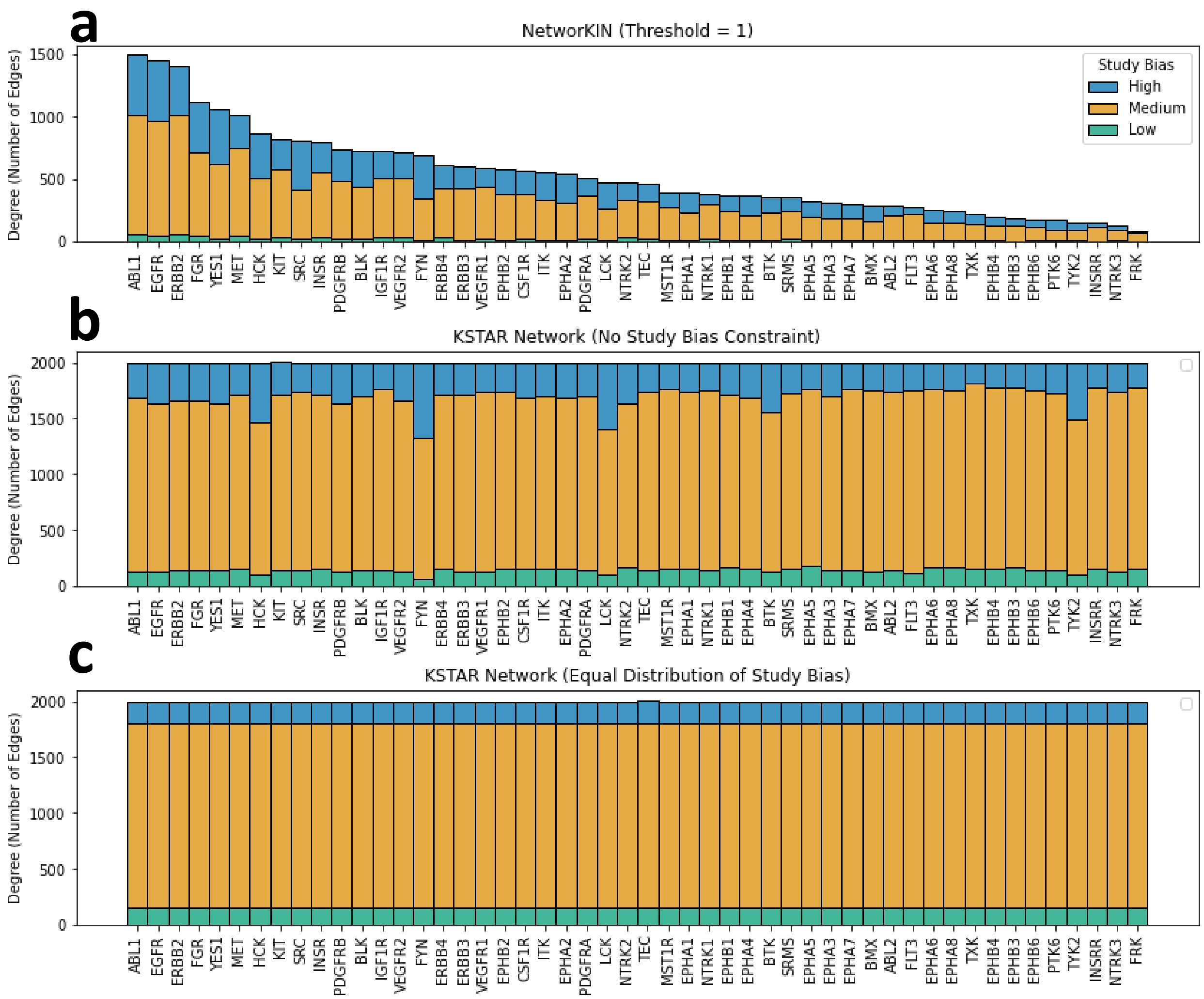


**Figure SM.3. KSTAR networks exhibit a more balanced distribution of kinase network edges** In order to account for kinase-specific study bias issues in kinase-substrate networks (certain kinases are connected to more substrates, or are connected to many well-studied sites), the heuristic prune enforces a rule that ensures all kinases have an equal number of edges in each pruned network, and that these edges have the same distribution of study bias for all kinases (as defined by the number of compendia each substrate is identified in). High study bias sites are found 3-5 compendia, medium study bias sites are found in 1-2 compendia, and low study bias sites are found in 0 compendia (only found in ProteomeScout). A) Number of substrates each kinase is connected to in NetworKIN thresholded with a value of 1, and the distribution of study bias within these connections. In addition to certain kinases having significantly more predicted substrates in these networks, we discovered that certain kinases, such as FYN and SRC tended to be connected to more well studied sites. B) Number of substrates each kinase is connected to in a pruned network if no study bias constraint is applied (Algorithm 1). While all kinases now have an equal number of edges, certain kinases tend to be more well connected to well studied sites (such as LCK, HCK, and FYN), leading to higher than expected, kinase-specific false positive rates. See Supplementary Figures 2 for examples. C) Number of substrates each kinase is connected to in one of the pruned KSTAR networks when the study bias constraint is applied (Algorithm 2). Unlike in B, all kinases have the same distribution of study bias among their substrate connections. This is an example of network used for the predictions in the main body of this work.

**Problem 6:** In thresholded kinase-substrate prediction networks, kinases from the same family commonly exhibit high substrate overlap, where many of the same phosphorylation sites provide evidence for the same kinases. This reduces the ability of prediction algorithms to discriminate between the activity of these kinases.

**Solution:** We have observed that the probability based selection of edges and the constraints placed on this selection (equal distribution of study bias, all kinases have the same number of edges, limits to the number of kinases a substrate can provide evidence for, etc.) during the heuristic prune procedure serve to significantly reduce the overlap between like kinases, allowing for greater discriminability in the activity of these kinases.

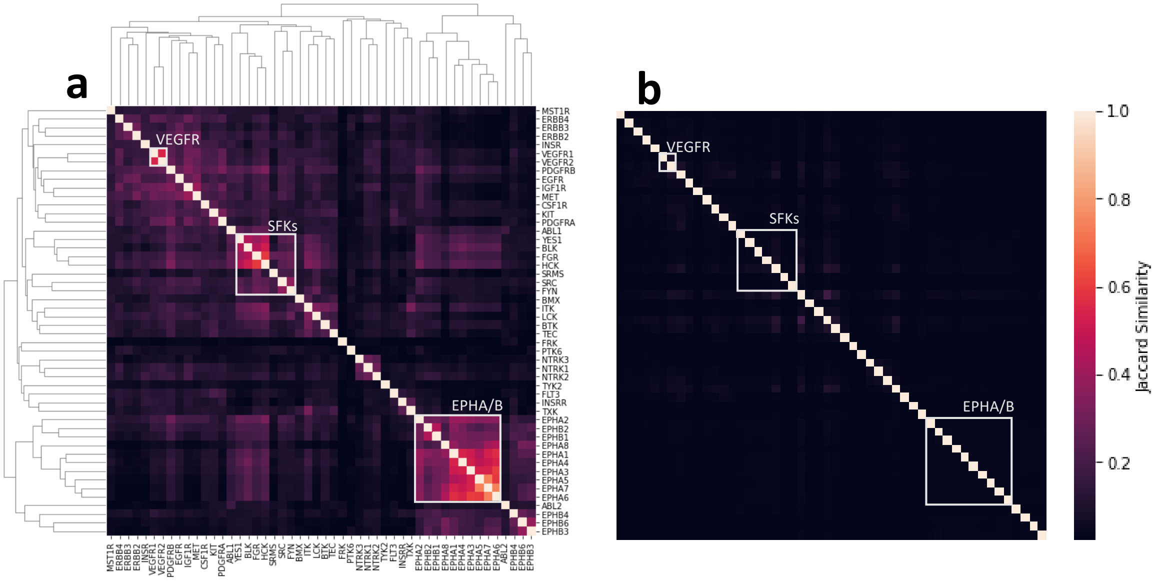


**Figure SM.4. Effect of Heuristic Prune on Network Similarity Between Kinases** A) Heatmap depicting the overlapping substrates between kinases in NetworKIN when thresholded by a value of 1, as defined by the Jaccard similarity index of the sites providing evidence for each kinase. Kinases were sorted using hierarchical clustering. Key kinases/kinase families that exhibit high overlap are indicated by the boxes. B) Heatmap depicting the overlapping substrates between the same kinases in KSTAR networks, as defined by the average Jaccard Similarity across all KSTAR networks. Kinases were sorted using the same order in A.

##### Challenges with Kinase Activity Inference

**Problem 7:** Quantification is required for the majority of currently available kinase activity inference algorithms. This is problematic for several reasons:

1. Relative quantification is inherently noisy, and a fold change can mean different things for different proteins. For example, a fold change of 2 can indicate an increase in abundance from 0.1 to 0.2, but it can also indicate an increase in abundance from 10 to 20, two very different outcomes.
2. In the clinical setting, it is often difficult to obtain a matched healthy sample required for relative quantification.

**Solution:** Measured abundance values are converted to binary evidence based on some cutoff value relevant to the biological problem, and binary evidence is used as input into the KSTAR algorithm. We can also convert to binary evidence based only on whether a site is identified in a sample. This also allows for use of the KSTAR algorithm without relying on any quantification at all, using any sites identified in a sample as evidence, as done for Figure 3 in the main text.

**Problem 8:** Experiments are more likely to identify well-studied sites, which can lead to experiment-specific false positive rates (see Supplementary Figures 2).

**Solution:** Compare enrichment p-values obtained from the real experiment to enrichment p-values that can be obtained from random phosphoproteomic experiments with the same properties as the real experiment (same number of sites used as evidence, same distribution of study bias). This is done using the Mann Whitney U-test, a non-parametric distribution test.

#### Implementing the KSTAR algorithm

In this section, we will expand upon our original description of the KSTAR algorithm and how it is implemented in practice. We have seperated the algorithm into two main sections for the purpose of clarity. First, we will describe the heuristic prune and how to generate the ensemble of possible kinase-substrate networks. For this section, the primary data that is required is a weighted kinase-substrate network containing predictions for the entire phosphoproteome, such as NetworKIN (Horn, 2014), GPS (Wang, Proteomics and Bioinformatics, 2020), or PhosphoPICK (Patrick, Biochimica et Biophysica Acta - Proteins and Proteomics, 2016), and the reference human phosphoproteome obtained from KinPred (Xue, PLOS Computational Biology, 2021). The second section describes how phosphoproteomic data is converted into kinase activities using the network ensemble obtained in the first section.

##### The Heuristic Prune and the Generation of the Network Ensemble

The heuristic prune generates an ensemble of binary kinase-substrate networks representing possible represenions of kinase-substrate interactions based on weighted graph predictions (we used NetworKIN, but theoretically can be any weighted kinase-substrate network). In this procedure, we ensure that all kinases have an equal number of edges/interactions in each binary network, set by *kinase_network_size*. We also ensured that no phosphorylation site provides evidence for more kinases than a predetermined cutoff, *site_limit*. Lastly, we generate seperate networks for serine/threonine sites and tyrosine sites, as these are nonoverlapping networks. Once these parameters have been determined, edges are randomly selected from the weighted network and added to the pruned network according to a probability equal to edge weights in the weighted network. Importantly, each kinase in the network will have edges added at the same rate (all kinases will have one edge in the pruned network before any have two, etc.), which ensures that every kinase is most likely to have their highest probability edges present in the final networks. If any substrate in the pruned network obtains a number of edges equal to *site_limit*, it is removed from the weighted network and cannot serve as evidence for any other kinases. Because edge selection is ultimately probabilistic in nature, every network generated using this procedure will be a unique representation of a possible kinase-substrate landscape. The implementation of this approach for a single network is described in Algorithm 1.

**Algorithm 1:** Heuristic prune of kinase-substrate network

**Data:** Weighted kinase-substrate network

**Input:** *kinase_network_size* = number of substrate interactions for each kinase in the pruned network

*site_limit* = maximum number of kinase interactions for each substrate in the pruned network

*modified_sites* = type of phosphorylation event to generate network for. Can either be Y (tyrosine) or ST (serine/threonine)

**Output:** Binary kinase-substrate network

```
Initialize empty binary network;
Reduce weighted network to include either only Y or only S/T sites;
for i = 1, 2, 3, ..., kinase_network_size do
    Shuffle kinase order;
    foreach kinase do
        substrate ← sample from weighted network with probability equal to kinase edge weights;
        Add substrate-kinase edge to binary network;
        Remove substrate-kinase edge from weighted network;
        if degree(substrate) >= site_limit then
            Remove substrate node from weighted network;
        end if
    end foreach
end for
return
```

We noticed that the above approach resulted in a high number of false positives for certain kinases and experiments (see Problem 5 and Supplementary Figures 2), and determined that this was likely a result of the study bias found within these networks, where well studied substrates tend to have high influence on predictions. To account for study bias, we modified the approach described in Algorithm 1 so that each kinase has edges/interactions with the same distribution of study bias (i.e. no kinases have only well-studied sites providing evidence for them, and no kinases have only poorly studied sites providing evidence for them). To do so, we defined study bias for each substrate in the human phosphoproteome as the number of different phosphoproteome compendia they are identified in. The compendia used in this study are PhosphoSitePlus (Hornbeck, Nucleic Acids Research, 2012), phosphoELM (Diella, BMC Bioinformatics, 2004), HRPD (Prasad, Nucleic Acids Research, 2009), dbPTM (Lu, Nucleic Acids Research, 2013), and UniProt (Uniprot Consortium, Nucleic Acids Research, 2021).

The heuristic prune described in Algorithm 2 is similar to Algorithm 1, except that each kinase in the final binary network contains an equal distribution of sites found in 0, 1, 2, 3, 4, or 5 compendia (Figure SM.3C). Algorithm 2 is the prune implementation we utilize throughout the main body of this work.

**Algorithm 2:** Heuristic prune of kinase-substrate network that ensures equal distribution of study bias for each kinase

**Data:** Weighted kinase-substrate network
Human Reference Phosphoproteome from KinPred
**Input:** *kinase_network_size* = number of substrate interactions for each kinase in the pruned network
*site_limit* = maximum number of kinase interactions for each substrate in the pruned network
**Output:** Binary kinase-substrate network

```
Initialize empty binary network;
Reduce weighted network to include only Y or S/T sites;
for n = 1, 2, 3, 4, 5 do
    /* In order to know how many sites found in n compendia to sample, we need to
       determine the fraction of these sites found in the overall phosphoproteome.
       Out of the total kinase_network_size, we will then sample that same fraction
       so that each kinase has the same distribution of study bias as the background
       network.                                                                  */
    compendia_size ← (number of sites in weighted network found in n compendia / total number of sites in weighted network) * kinase_network_size;
    compendia_network ← reduce weighted network to only include sites found in n compendia;
    /* For each compendia size, n, follow the same pruning procedure as described in
       Algorithm 1.                                                              */
    for i = 1, 2, 3, ..., compendia_size do
        Shuffle kinase order;
        foreach kinase do
            substrate ← sample from compendia_network with probability equal to kinase edge
                    weights;
            Add substrate-kinase edge to binary network;
            Remove substrate-kinase edge from weighted network;
            if degree(substrate) >= site_limit then
                Remove substrate node from weighted network;
            end if
        end foreach
    end for
end for
```

##### Calculating Kinase Activity from a Phosphoproteomic Experiment

Once networks have been generated, we are now ready to predict kinase activity based on the phosphorylation sites identified in a mass spectrometry experiment. Prior to enrichment calculations, the list of phosphorylation sites to use as evidence for a particular sample must be determined. In most cases, this is determined by defining *threshold* and accepting any phosphorylation sites with abundance values greater than the *threshold* as evidence. In other cases, we either do not want to use the relative abundance values or do not have relative abundance values (no matched samples), and instead want to use all sites identified in an experiment as evidence (in practice, this equivalent to setting a very low *threshold* like -1e10).

The first step of activity prediction is to assess the statistical enrichment of each kinase's substrates identified in a sample using the hypergeometric distribution. In this test, for each kinase, we are asking what is the likelihood that there are $k$ substrates of a kinase in the the $n$ phosphorylation sites providing evidence for a sample, based on the total number of substrates ($K$) and phosphorylation sites ($N$) in the background phosphoproteome. As there are a total of 50 binary networks in the ensemble, the end result of this process is to produce 50 p-values for each kinase. The implementation of this procedure is described in Algorithm 3.

**Algorithm 3:** Calculating statistical enrichment of kinase substrates in binary networks

```
Data: Phosphoproteomic experiment
      Ensemble of pruned networks generated from Algorithm 2
Input: threshold = cutoff value that determines which sites are used as evidence
Output: Arrays with 50 p-values (one for each kinase) containing hypergeometric enrichment
        results in each of the 50 pruned networks

Intialize p-value array;
binary experiment ← reduced experiment including phosphorylation sites with abundance
        >= threshold;
foreach kinase do
    foreach network do
        k ← number of kinase substrates in binary experiment;
        K ← number of kinase substrates in network;
        n ← number of sites in binary experiment;
        N ← number of sites in network;
        pval ← 1 − hypergeometric_cdf(k − 1, K, n, N);
        Add pval to array of p-values;
    end foreach
end foreach
```

Next, we wished to ask whether, for each kinase, the distribution of p-values obtained from Algorithm 3 could be obtained from a random experiment where substrates are randomly pulled from the human phosphoproteome. To do so, we first needed to generate a collection of random experiments. In order to accurately reflect the characteristics of the real dataset, each random experiment must have the same number of observed phosphorylation sites and equal distribution of study bias (defined by the number of compendia a site is identified in). The procedure of generating these random experiments is described in Algorithm 4.

**Algorithm 4:** Generate random phosphoproteomic experiments for use in Mann Whitney tests

```
Data: Phosphoproteomic experiment (Real)
      Human Phosphoproteome Compendia (KinPred)
Input: num_experiments = number of random experiments to generate
Output: random_array = array of random experiments with identical characteristics as the real
        experiment (number of phosphorylation sites, distribution of study bias, etc.).

Initialize random_array;
for i = 1, 2, 3, ..., num_experiments do
    Initialize random_experiment;
    foreach site identified in real experiment do
        numCompendia ← number of compendia site is found in;
        random_site ← randomly sampled site from human phosphoproteome compendia found in
                the same number of compendia as numCompendia;
        Add random_site to random_experiment
    end foreach
    Add random_experiment to random_array
end for
```

Once random experiments have been generated in Algorithm 4, we can generate the final activity scores for each kinase. Ultimately, this score is a reflection of the enrichment of kinase-substrates in the actual experiment (calculated in Algorithm 3), relative the enrichment that could be obtained from a random phosphoproteomic experiment. For this task, we use a Mann Whitney U-test, non-parametric distribution-based test which compares the ranked p-values of the real and random experiments. The outcome of this test is a p-value for each kinase, which we then convert to an activity score by taking the -log10 of each p-value. In this way, kinases with high activity (and as a result of high enrichment of substrates in the dataset) will have high activity scores. The application of this test is demonstrated in Algorithm 5.

**Algorithm 5:** Calculate the final KSTAR activity score using a Mann Whitney U test for an individual kinase

**Input:** *real_p* = array of 50 p-values obtained from applying Algorithm 3
*random_array* = 150 random experiments generated from Algorithm 4
**Output:** *activity* = -log10(MannWhitney p-value)

*random_p* ← Algorithm 3 applied to each random experiment in *random_array*;
**foreach** *kinase* **do**
    $MannWhitneyP \leftarrow MannWhitneyUtest(real\_p_{kinase}, random\_p_{kinase})$;
    $activity_{kinase} \leftarrow -log10(MannWhitneyP)$;
**end foreach**
**return** activity;

To verify the significance of these scores, we then ask how often we could obtain the same activity scores or better from the random datasets. To do so, we treat one random experiment generated from Algorithm 4 as the real experiment and repeat Algorithm 5 to generate a random activity score. We then repeat this procedure to generate an array of random activity scores. The false positive rate is then the fraction of times a random experiment generated an activity score equal to or greater than the activity score calculated for the real dataset.

**Algorithm 6:** Calculate the false positive rate of obtaining the activity score for a single kinase obtained from Algorithm 5

**Input:** *real_activity* = activity score obtained from Algorithm 5
*random_array* = 150 random experiments generated from Algorithm 4
*numTrials* = number of random activity scores to obtain for FPR calculation
**Output:** *fpr* = fraction of random experiments which obtained the same or greater activity score as *real_activity*

Initialize *numFalsePositives* (*numFalsePositives* = 0);
**for** $i = 1, 2, 3, ..., numTrials$ **do**
    /* randomly select one of the random experiments in *random_array* to be treated as a real experiment in the MannWhitneyU test */
    *pseudoreal* ← randomly select experiment from *random_array*;
    *trimrandom* ← *random_array* with *pseudoreal* removed;
    /* So, *pseudoreal* should consist of 1 experiment from *random_array*, and *trimrandom* should consist of all other experiments from *random_array* */
    *MannWhitneyP* ← apply algorithm 5, treating *pseudoreal* as the real experiment and *trimrandom* as random experiments;
    $random\_activity_i \leftarrow -log10(MannWhitneyP)$;
    **if** $random\_activity_i >= real\_activity$ **then**
        $numFalsePositives \leftarrow numFalsePositives + 1$
    **end if**
**end for**
$fpr \leftarrow \frac{numFalsePositives}{numTrials}$;
**return** *fpr*;
