## Supplemental Figure 1 for "KSTAR: An algorithm to predict patient-specific kinase activities from phosphoproteomic data"

### Supplementary Figures 1: Assessing the impact of pruning and parameter selection in KSTAR

#### Goal

1. Profile the impact of the heuristic prune on the characteristics of the kinase-substrate network
2. Explore the impact of applying different cutoffs to experimental datasets when determining which phosphorylation sites to use as evidence.

#### Methods

For comparison to commonly used kinase-substrate networks, kinase-substrate annotations were downloaded from PhosphoSitePlus. The full NetworKIN prediction graph was downloaded for the entire phosphoproteome and edges with weights less than 1 were removed. A threshold of 1 was selected in order to balance the number of edges that are removed from the network and the inaccuracy often observed at low thresholds.

To look at sensitivity of KSTAR results to different evidence cutoffs in phosphoproteomic experiments, we generated kinase activity predictions using different evidence sizes (number of phosphorylation sites identified in original experiment used for prediction) for a tyrosine dataset (K562 cells, (Di Palma et al., Journal of Proteomics, 2013)) and a serine/threonine dataset (BT-474 cells, (Wiechmann et al., ACS Chemical Biology, 2021)). In both cases, the first test used all of the sites identified in the experiment, and subsequent tests with smaller evidence sizes removed the least abundant sites from analysis to mimic the effect of thresholding.

#### Table of Contents

**S1.1** - Comparing the characteristics of pruned networks to PhosphoSitePlus and NetworKIN (Page 2)
**S1.2** - Assessing the stability of tyrosine predictions using different amounts of sites as evidence, mimicking the effect of applying a threshold to the phosphoproteomic data to select phosphosites used as evidences (Page 3)
**S1.3** - Assessing the stability of serine/threonine predictions using different amounts of sites as evidence, mimicking the effect of applying a threshold to the phosphoproteomic data to select phosphosites used as evidence (Page 4)

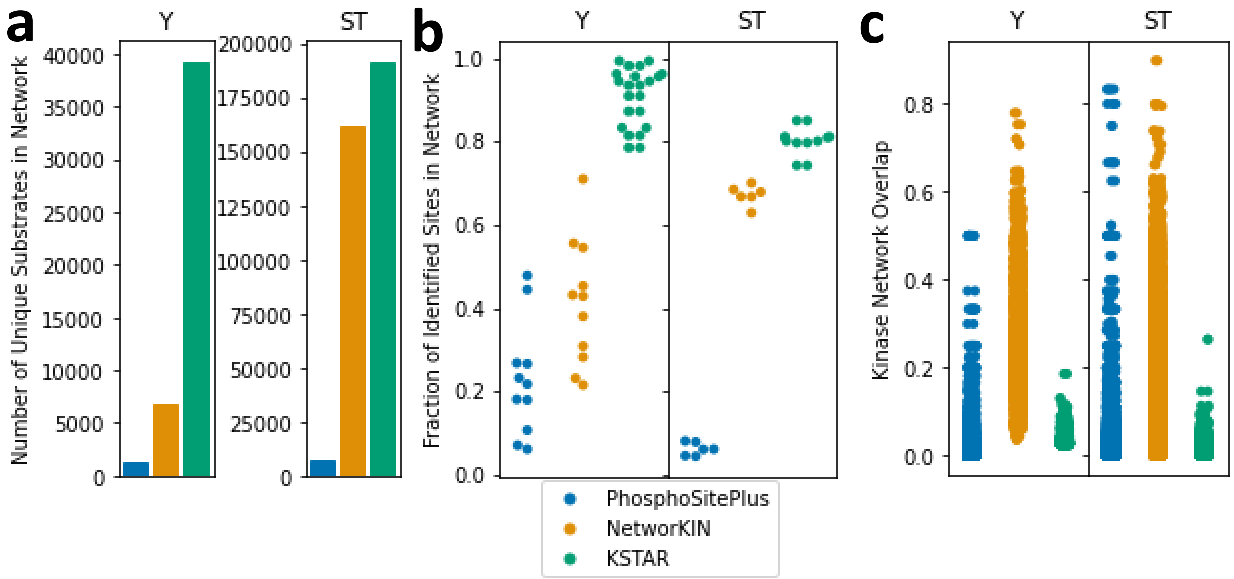


**Figure S1.1. Comparison of KSTAR pruned networks to PhosphoSitePlus and NetworKIN** Comparison of network characteristics for known kinase-substrate annotations from PhosphoSitePlus (blue), kinase-substrate predictions from NetworKIN thresholded with a value of 1 (remove edges with edge weights less than 1)(orange), and the KSTAR ensemble of pruned networks (green). A) Number of unique substrates found within each network. B) Fraction of sites identified within an experiment that are also found within the kinase-substrate network. Experiments include any phosphoproteomic dataset for which KSTAR predictions were applied (Predictions found in Figures 2-4), a total of 11 experiments. Each point on the plot represents a unique experiment. While prediction algorithms using either PhosphoSitePlus or NetworKIN will have to drop $> 50\%$ of sites identified in an experiment, KSTAR typically loses $< 20\%$. This problem is especially apparent in tyrosine networks. C) Pairwise network similarity between different kinases in each network, defined by the Jaccard similarity, calculated as $\frac{total\ number\ of\ substrates\ connected\ to\ both\ kinases}{total\ number\ of\ substrates\ of\ connected\ to\ either\ kinase}$. Each point represents a different pairwise comparison. Unlike PhosphoSitePlus and NetworKIN, the majority of kinases share $< 20\%$ of substrates in KSTAR networks.

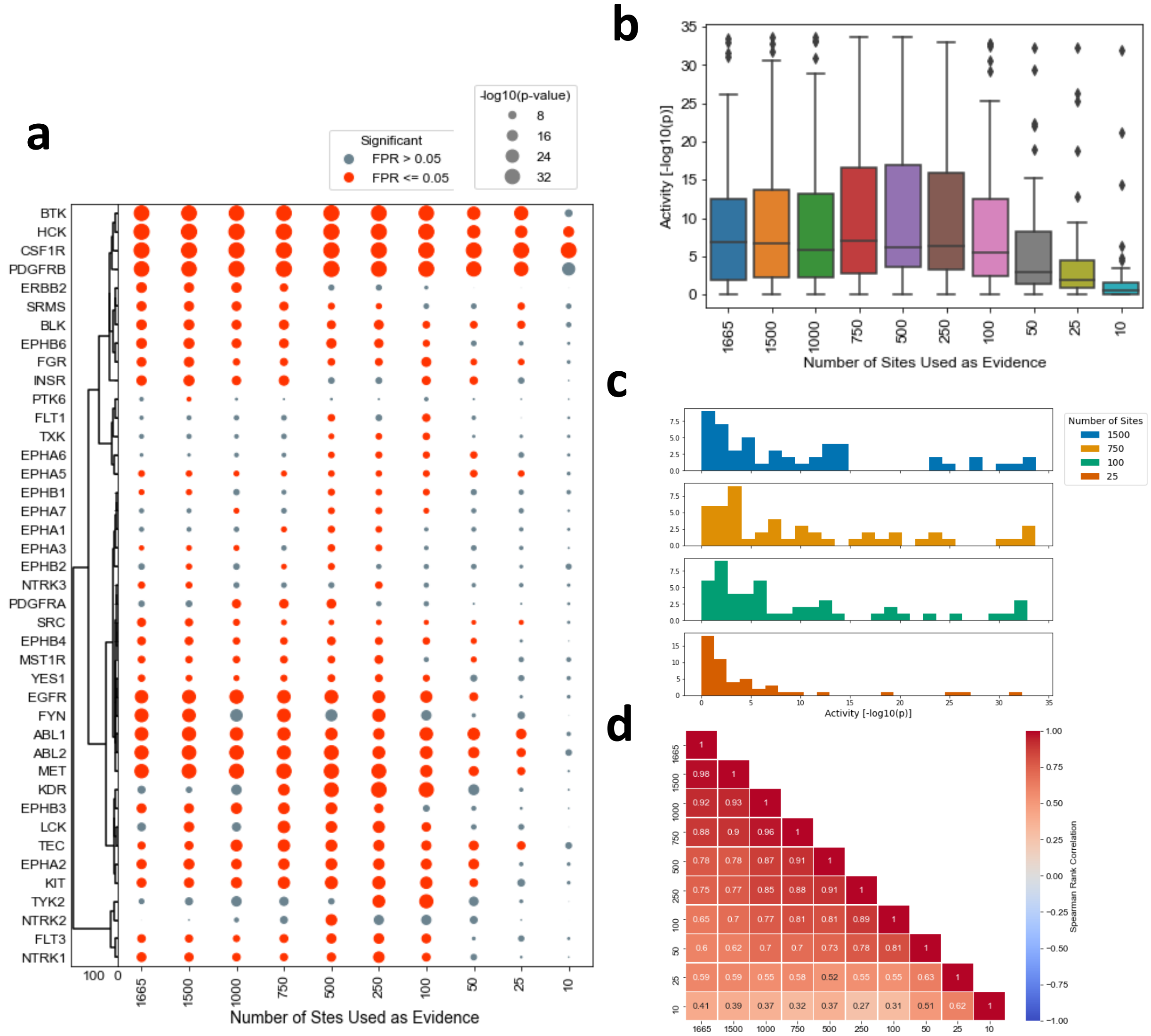


**Figure S1.2. Impact of experiment threshold on tyrosine kinase predictions** Using control data of K562 chronic myeloid leukemia cell lines (Di Palma et al., Journal of Proteomics, 2013), we assessed variance in KSTAR predictions for tyrosine kinases using different thresholds (cutoff value to deterimine whether a phosphorylation site should be included in evidence). The threshold determines the total number of sites used as evidence for prediction. For this test, predictions were based evidence sizes ranging from 10 - 1665 (all identified) sites. A) KSTAR predictions for kinases that have predicted activity in at least one test ($FPR <= 0.05$). B) Distribution of KSTAR activity scores for all kinases at each evidence size. C) Histograms demonstrating the distribution of activity scores when 25, 100, 750, 1500 sites are used for prediction. D) Correlation of activity rankings between predictions of different evidence sizes.

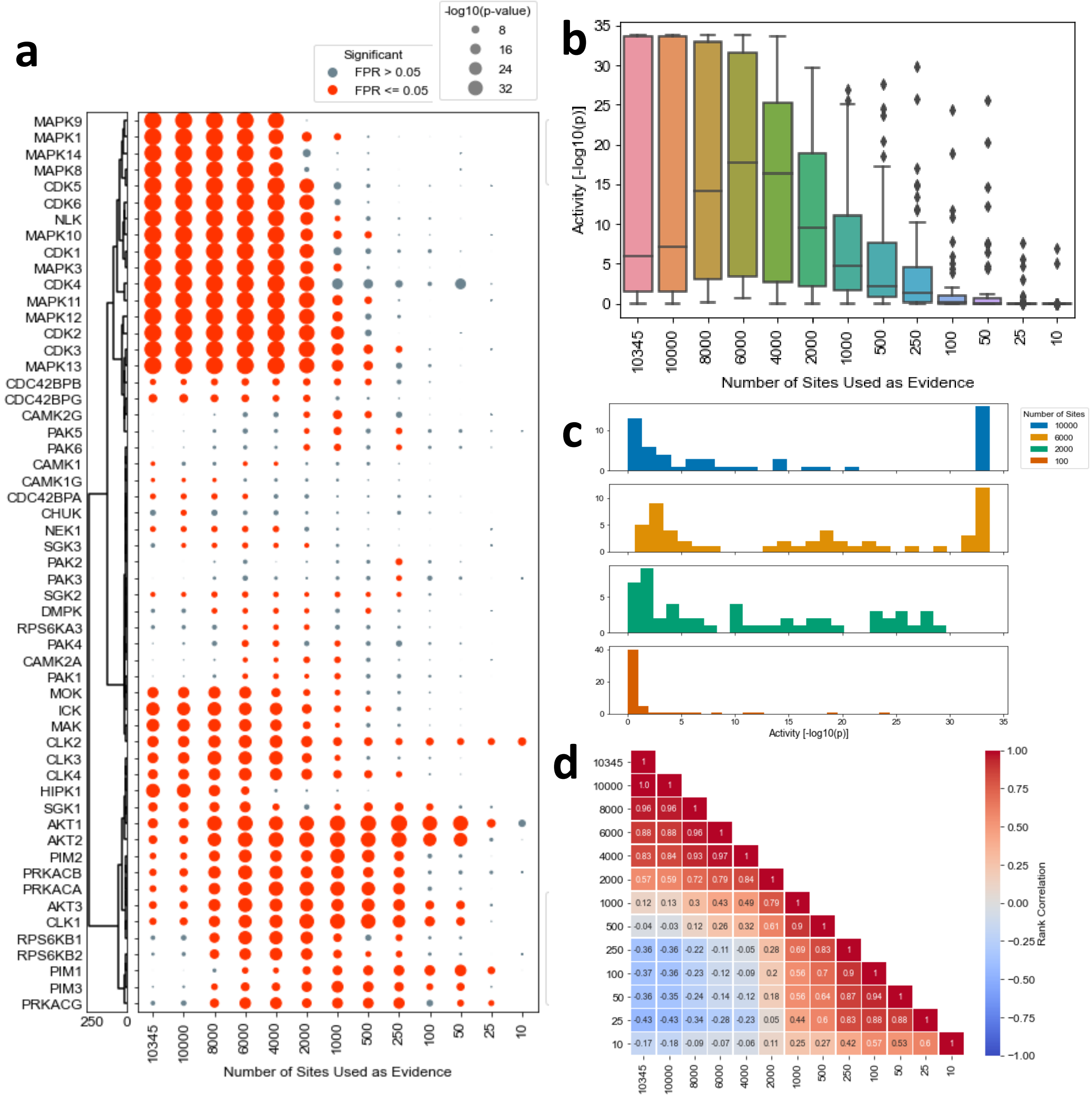

**Figure S1.3. Impact of experiment threshold on serine/threonine kinase predictions** Using control data of BT-474 Breast cancer cell lines (Wiechmann et al., ACS Chemical Biology, 2021), we assessed variance in KSTAR predictions for serine/threonine kinases using different thresholds (cutoff value to deterimine whether a phosphorylation site should be included in evidence). The threshold determines the total number of sites used as evidence for prediction. For this test, predictions were based evidence sizes ranging from 10 - 10345 (all identified) sites. A) KSTAR predictions for kinases that have predicted activity in at least one test ($FPR <= 0.05$). B) Distribution of KSTAR activity scores for all kinases at each evidence size. C) Histograms demonstrating the distribution of activity scores when 100, 2000, 6000, or 10000 sites are used for prediction. D) Correlation of activity rankings between predictions of different evidence sizes.
