## Supplemental Figure 2 for "KSTAR: An algorithm to predict patient-specific kinase activities from phosphoproteomic data"

### Supplementary Figures 2: Controlling for kinase- and experiment-specific false positive rates

#### Goal

1. Assess the bias found within different phosphoproteomic databases by predicting activity from random samplings of the database
2. Assess the bias found with phosphoproteomic datasets by looking at the distribution of study bias across the sites identified in an experiment
3. Assess the relationship between study bias of a substrate and quantified log fold changes

#### Methods

In order to measure the false positive rate, we randomly created 200 random datasets, each composed of 250 randomly selected phosphotyrosine sites using different phosphoproteomic databases as the background. We set the desired false positive rate at 0.05 and therefore expect, on average, about 10 positives per kinase across all datasets. The false positive results are from KSTAR networks before the final addition of controlling for the distribution of compendia was added (Figures S2.1 - S2.5). The size of the dataset was selected based on common dataset sizes for phosphotyrosines and this value is much smaller than the smallest of the compendia background.

#### Summary of Results

The first four figures here have been ordered from backgrounds that produce the highest false positive rates to those that produce the lowest false positive rates. It became clear that the kinases with high false positive rates were consistent across all compendia (except those from PhosphoSitePlus, which is the largest and predominantly comprised of mass spectrometry identified sites). In short, FYN, LCK, and HCK demonstrated significant levels (100%) false positive rates. On the other end of the range, it became clear that most kinases were producing less than expected false positive rates (0%), which suggested that it is also more difficult to yield true positives for these kinases. Additionally, we noticed a trend that the false positive rates were highest from the compendia that overlapped most with the genesis of the NetworKIN prediction models (phosphoELM, S2.1) and lowest from the compendia most separated from training data that formed the networks (PhosphoSitePlus, S2.4). We therefore hypothesized that there was a direct connection between the study bias of the phosphorylation sites (the more compendia a site is in the more likely it is annotated and was used in training the networks). Figure S2.5 shows that our original KSTAR networks that generated such kinase-specific false positive rates indeed showed skew for more studied substrates connected to the kinases yielding high false positive rates. We next asked if we could measure the false positive rate for random experiments by random draws from the human phosphoproteome. We found that all experiments analyzed in this manuscript were not representative of random samples from the phosphoproteome, with respect to how many compendia substrates were annotated by (Fig. S2.7). We also found that magnitude of fold changes (i.e. quantification) was partially related to the study bias of the individual substrate (Fig. S2.8). This suggests that each experiment will have different false positive rates, influenced directly by how well studied the sites are within the dataset. This motivated the final mitigation strategy of selecting random datasets for measuring empirical false positive rates based on sampling sites from the phosphoproteome, such that the distribution matches that of the real experiment. The combination of avoiding quantification, normalizing kinase study bias distribution (Fig. S2.6), and sampling random sets to control for experiment-specific distributions, helps to control the false positive rate as desired in a kinase- and experiment-specific manner.

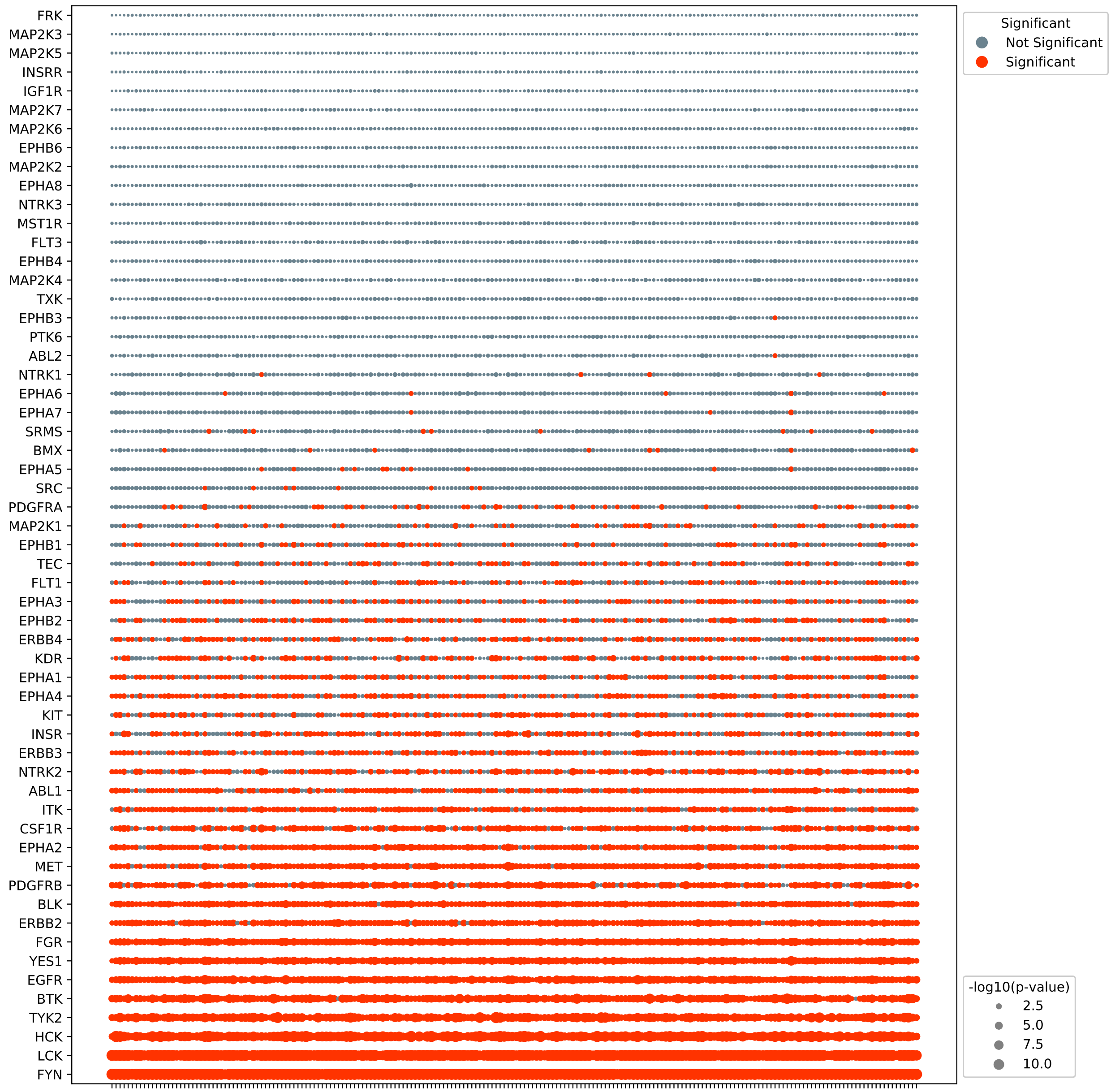


**Figure S2.1. Kinase-Specific False Positive Rates in PhosphoELM** To assess false positive rates obtained from PhosphoELM, 200 random datasets were created by randomly sampling 250 phosphotyrosine sites from PhosphoELM, and KSTAR was applied (without accounting for study bias, i.e. no study bias constraint in heuristic prune or comparison to random activities from Mann Whitney U test) to generate activity predictions for each random dataset. Each column in the above dotplot corresponds to results from a different random datset. The size of each dot corresponds to the median p-value enrichment obtained across all pruned networks, and is colored based on significance ($p <= 0.05$). Many kinases, including FYN, LCK, and HCK exhibited close to a 100% false positive rate, while others like FRK and INSRR were not predicted active in a single random dataset, lower than the expected 5% false positive rate.

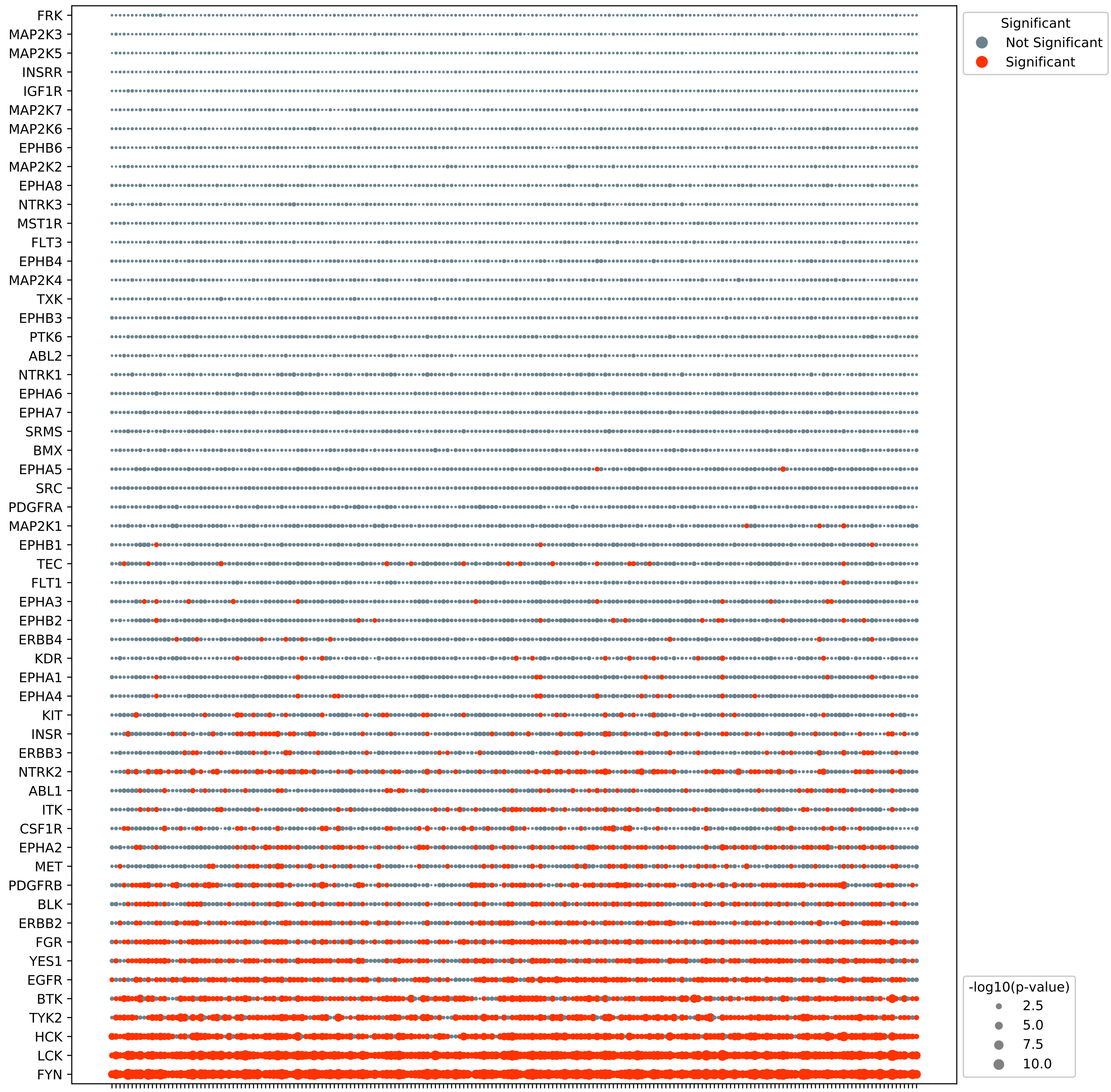


**Figure S2.2. Kinase-Specific False Positive Rates in HRPD** To assess false positive rates obtained from HPRD, 200 random datasets were created by randomly sampling 250 phosphotyrosine sites from HPRD, and KSTAR was applied (without accounting for study bias, i.e. no study bias constraint in heuristic prune or comparison to random activities from Mann Whitney U test) to generate activity predictions for each random dataset. Each column in the above dotplot corresponds to results from a different random datset. The size of each dot corresponds to the median p-value enrichment obtained across all pruned networks, and is colored based on significance ($p <= 0.05$). Similar to Figure S2.1, many kinases, including FYN, LCK, and HCK exhibited close to a 100% false positive rate, while others like FRK and INSRR were not predicted active in a single random dataset, lower than the expected 5% false positive rate.

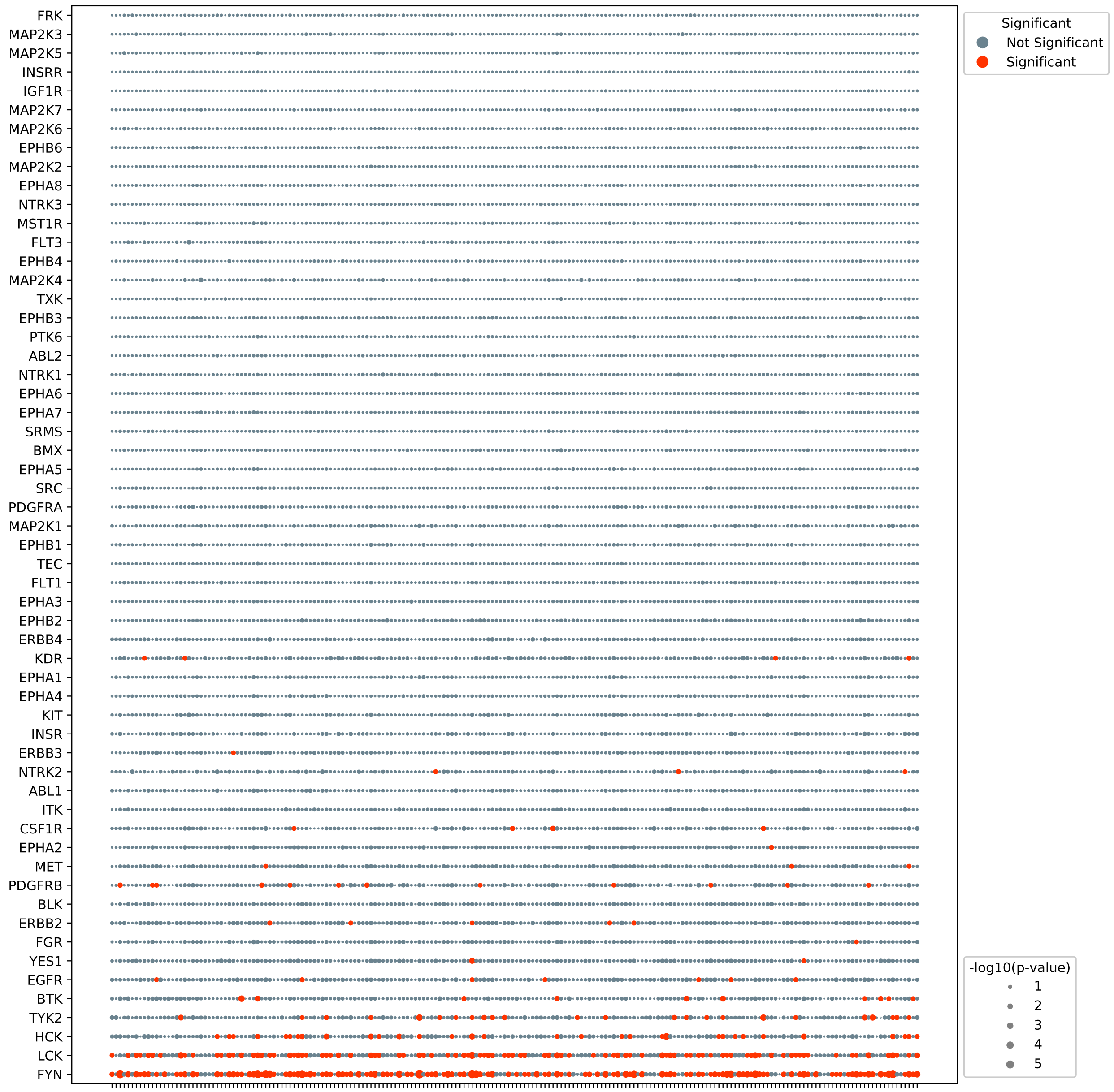


**Figure S2.3. Kinase-Specific False Positive Rates in dbPTM** To assess false positive rates obtained from dbPTM, 200 random datasets were created by randomly sampling 250 phosphotyrosine sites from dbPTM, and KSTAR was applied (without accounting for study bias, i.e. no study bias constraint in heuristic prune or comparison to random activities from Mann Whitney U test) to generate activity predictions for each random dataset. Each column in the above dotplot corresponds to results from a different random datset. The size of each dot corresponds to the median p-value enrichment obtained across all pruned networks, and is colored based on significance ($p <= 0.05$). Predictions exhibit lower false positve rates than when sampling from HPRD or PhosphoELM, but still see high kinase-specific false-positive rates for kinases like FYN, LCK, and HCK, and sees more kinases with 0 positives across all datasets.

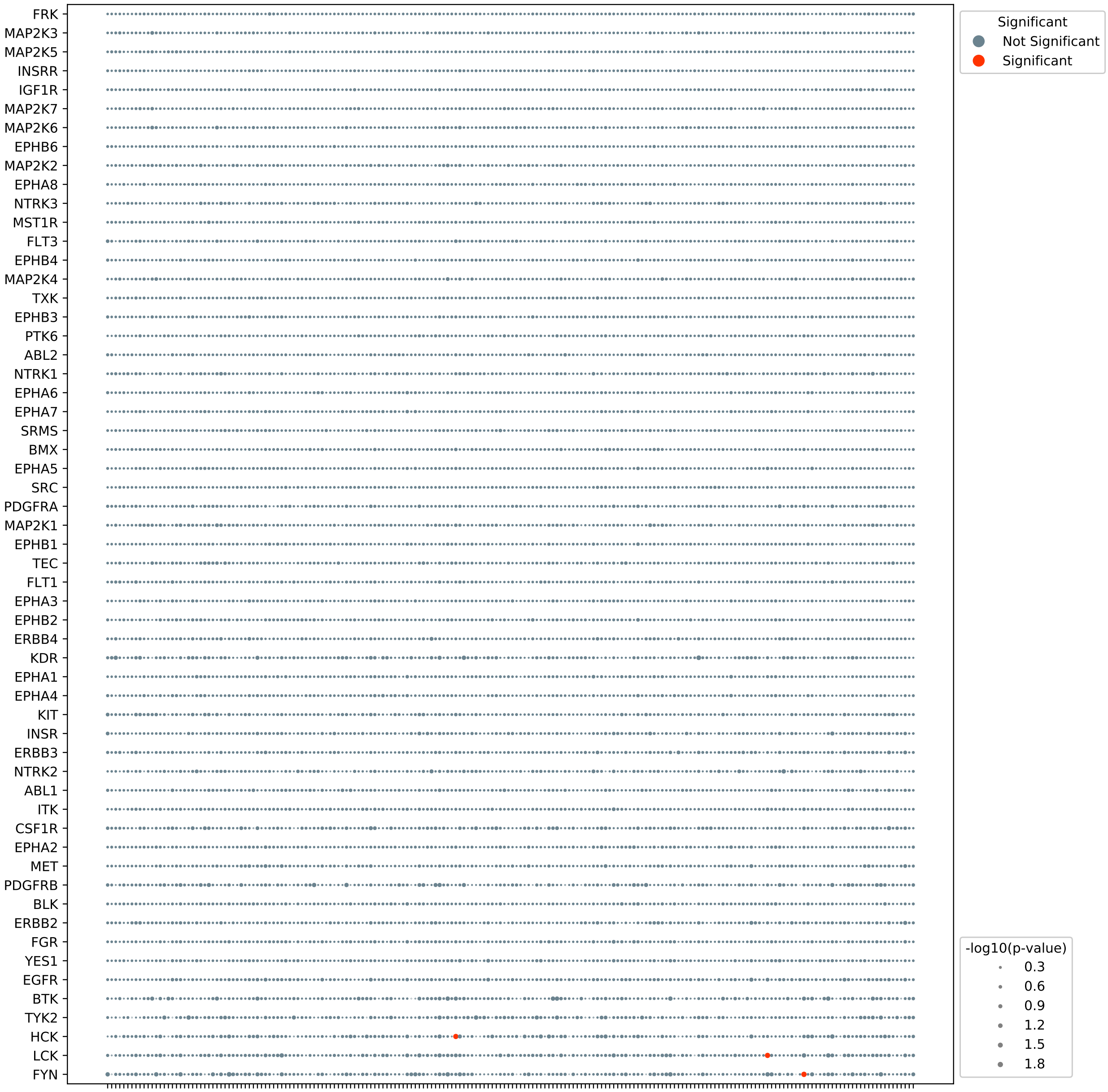


**Figure S2.4. Kinase-Specific False Positive Rates in PhosphoSitePlus** To assess false positive rates obtained from PhosphoSitePlus, 200 random datasets were created by randomly sampling 250 phosphotyrosine sites from PhosphoSitePlus, and KSTAR was applied (without accounting for study bias, i.e. no study bias constraint in heuristic prune or comparison to random activities from Mann Whitney U test) to generate activity predictions for each random dataset. Each column in the above dotplot corresponds to results from a different random datset. The size of each dot corresponds to the median p-value enrichment obtained across all pruned networks, and is colored based on significance ($p <= 0.05$). All kinases show lower than expected false positive rates (0 instead of 10 positives) when sampled from PhosphoSitePlus as the background. PhosphoSitePlus is the largest of the compendia and derived predominantly of mass spectrometry identified sites from within the Cell Signaling Technology pipelines.

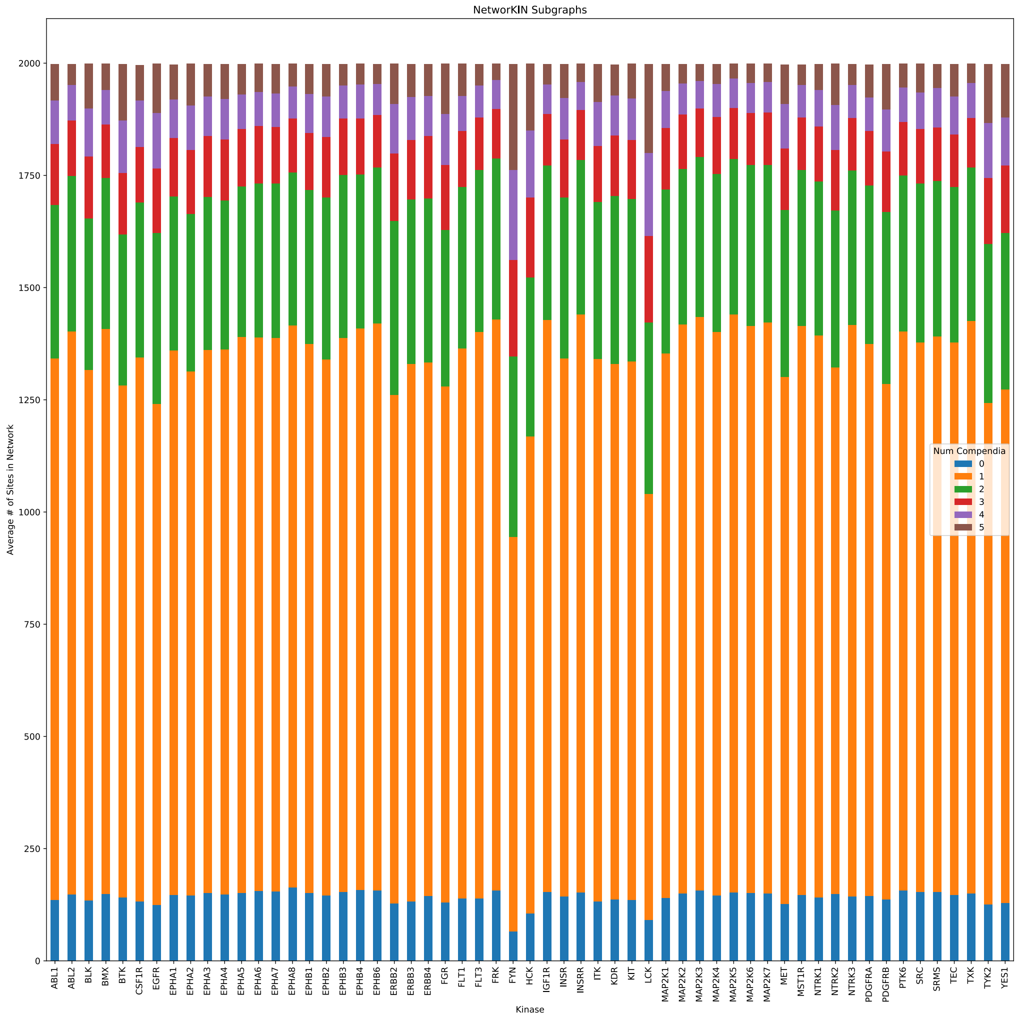


**Figure S2.5. Distribution of substrate study bias in original networks** We plotted distribution of substrates, based on the number of compendia they are observed in, for our original KSTAR generated networks (no study bias constraint, Algorithm 1 in supplemental methods) and found that the high false positive rates of kinases correlated with kinases highly connected to well studied phosphorylation sites. For example, FYN, LCK, and HCK have the highest proportion of phosphorylation sites from three or more compendia and are the kinases with the highest degree of false positives. The above plot represents the average number of substrates found in each study bias grouping (found in 0, 1, 2, 3, 4, or 5 compendia) across all of the 50 generated networks.

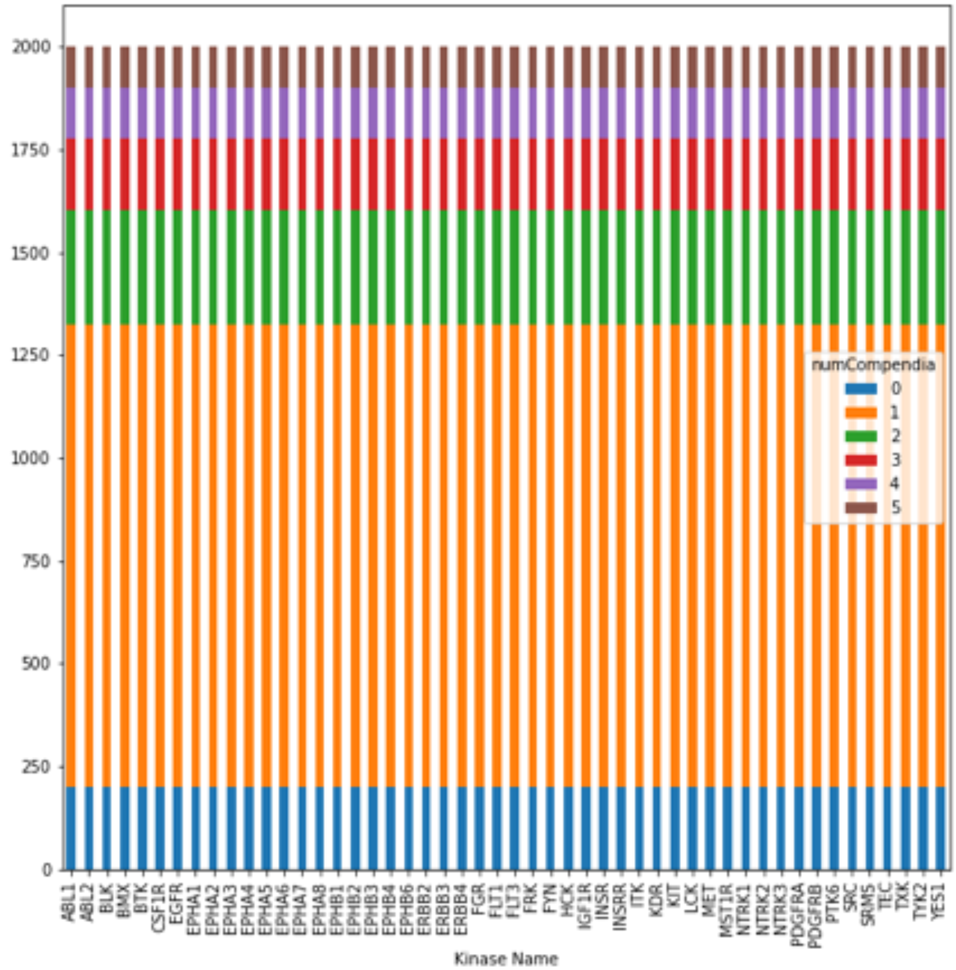


**Figure S2.6. Distribution of study bias in final networks** We added the constraint that all kinases should be connected to substrates with an equal distribution of study bias, as defined by the number compendia they are observed in. This ensures that no kinase will be connected to more well studied sites than any other kinase, which leads to kinase-specific false positive rates. This leads to a more balanced network, as observed in the above plot indicating the average number of substrates found in each study bias grouping (found in 0, 1, 2, 3, 4, or 5 compendia) across the final KSTAR generated networks (with a study bias constraint, Algorithm 2 in supplemental methods).

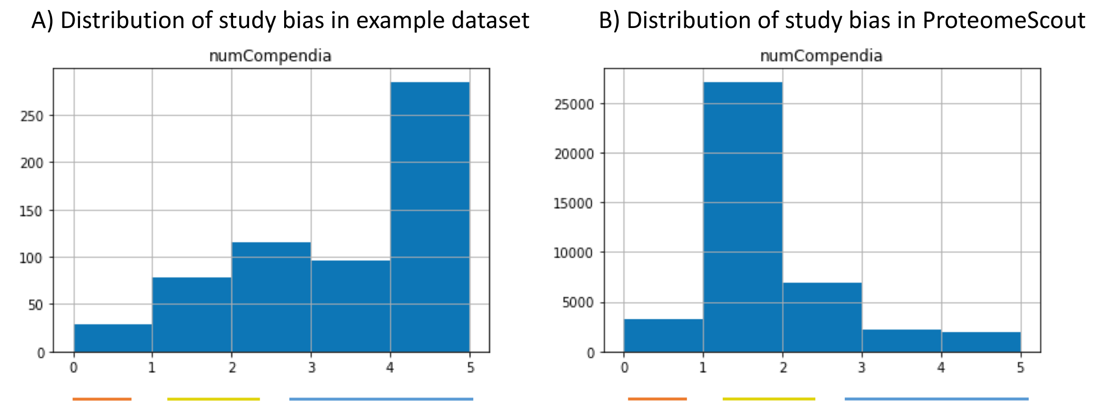


**Figure S2.7. Distribution of study bias in a real dataset, compared to whole phosphoproteome** We found that the phosphorylation sites from experiments were much more likely to contain sites that are well annotated than expected by random chance (i.e. contributing to experiment-specific study bias). **A)** This is the histogram of the number of compendia phosphorylation sites are observed in for the phosphotyrosines of the PDX dataset from Huang et al. (Nature Communications, 2017). **B)** This is the distribution for the entirety of the human phosphoproteome, which shows the majority of sites would be annotated by only one compendia (most likely, these are annotated only in PhosphoSitePlus). The three lines represent our final classes of study bias for low (0 compendia), medium (1-2 compendia), and high (more than 3 compendia). This classification was defined and published in our prior work (KinPred Xue et al., PLoS Computational Biology, 2021).

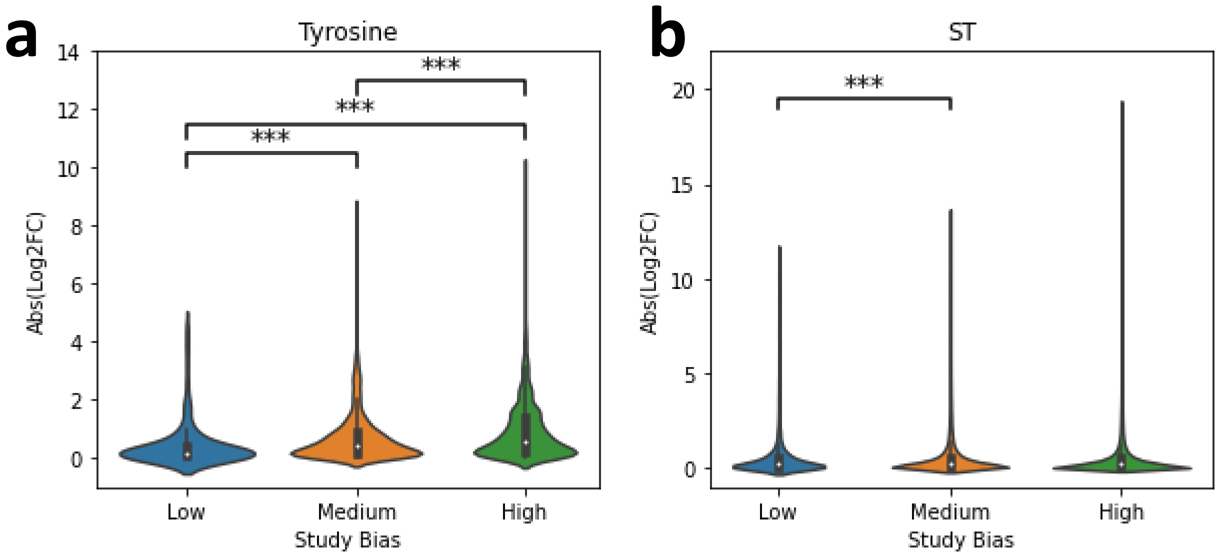


**Figure S2.8. Relationship between study bias and quantification in the benchmarking dataset** In addition to the likelihood of identification discussed in S2.7, we found that well studied tyrosine sites identified in an experiment tend to exhibit larger log2 fold changes than less well studied sites, according to a one-tailed Mann Whitney U test. We did not find the same relationship between study bias and quantification in serine/threonine datasets. To obtain these results, we Quantification values were obtained from all datasets in the benchmarking dataset used in Figure 3 of the main text and Supplemental Figure 4. Statistical signifance of log2 fold changes were calculated using a one-tailed Mann Whitney U-test and is indicated on the plot (*: $p \leq 0.01$, **: $p \leq 0.001$, ***: $p \leq 0.0001$). We have defined three classes of study bias based on the number of compendia a site is identified in: low (0 compendia), medium (1-2 compendia), and high (more than 3 compendia). This classification was defined and published in our prior work (KinPred, Xue et al., PLoS Computational Biology, 2021).
