## Supplemental Figure 3 for "KSTAR: An algorithm to predict patient-specific kinase activities from phosphoproteomic data"

### Supplementary Figures 3: Full KSTAR Predictions on Control Datasets

#### Goal

Provide the full KSTAR predictions on activation and inhibition datasets discussed in Figure 2

#### Methods

For all inhibition and activation datasets where KSTAR was applied (Figure 2), a dotplot is provided that includes all kinases with predictions (limited by the kinases with substrate predictions in NetworKIN). For the two serine/threonine datasets (Figure S3.5, S3.6), only kinases with significant activity in at least one sample ($FPR <= 0.05$) were included in the plot due to space constraints.

#### Table of Contents

**S3.1** - Full activity predictions during EGF stimulation of 184A1 epithelial cells (Page 2)
**S3.2** - Full activity predictions during EGF/HRG stimulation of 184A1 epithelial cells overexpressing HER2, corresponding to Figure 2A (Page 3)
**S3.3** - Full activity predictions during TCR activation of Jurkat cells, corresponding to Figure 2B (Page 4)
**S3.4** - Full activity predictions during BCR-ABL inhibition by dasatanib in chronic myeloid leukemia cells, corresponding to Figure 2C (Page 5)
**S3.5** - Full activity predictions after AKT Inhibition in breast cancer cells with 5 different AKT inhibitors, corresponding to Figure 2D (Page 6)
**S3.6** - Full activity predictions after BRAF inhibition in two different colorectal cancer cell lines leading to paradoxical MAPK activation, corresponding to Figure 2E (Page 7)
**S3.7** - Quantile-normalized activity scores for MAPK1 and MAPK3 after BRAF inhibition in two different colorectal cancer cell lines, corresponding to Figure 2E (Page 8)

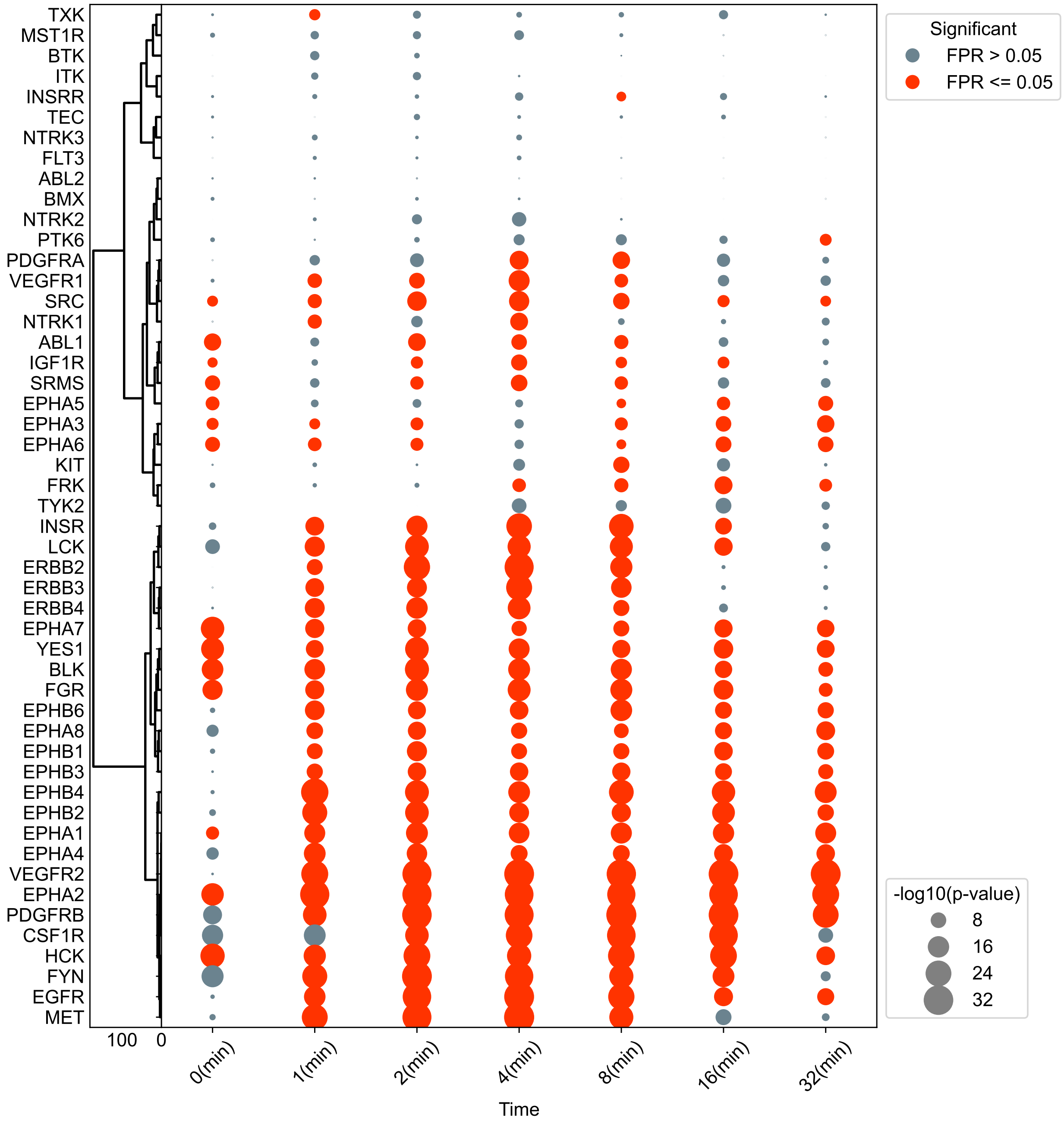


**Figure S3.1. EGF stimulation of 184A1 epithelial cells** Full KSTAR predictions on EGF stimulation phosphoproteomic data obtained from (Wolf-Yadlin et al., Proceedings of the National Academy of Sciences, 2007). 184A1 epithelial cells were stimulated with EGF and phosphorylation was measured at 0, 1, 2, 4, 8, 16 and 32 minutes. For each condition, sites with abundance ratios greater than 1 were used as evidence, where ratios were relative to the 4 minute timepoint. Kinases were sorted using hierarchical clustering with Ward linkages.

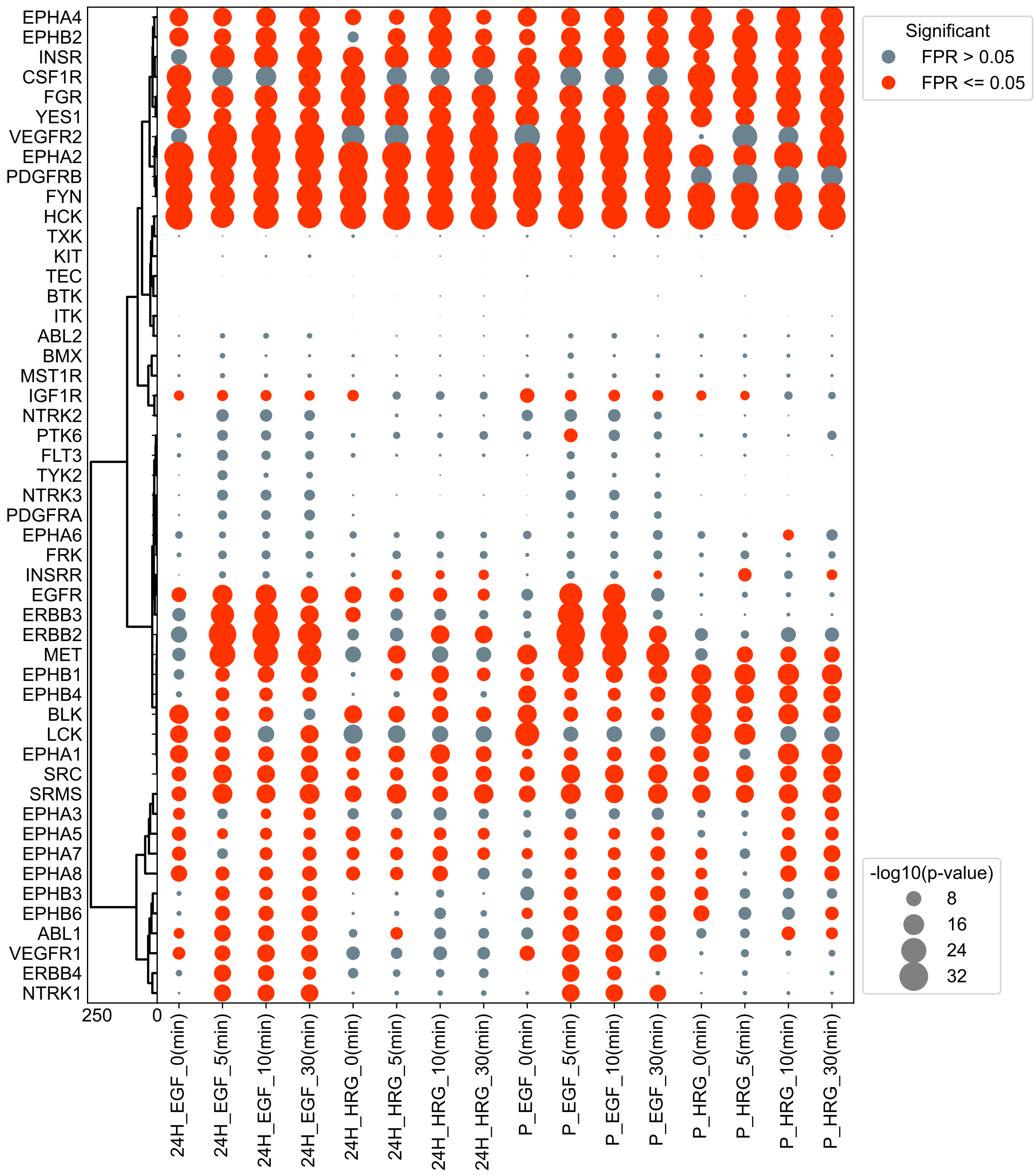


**Figure S3.2. EGF/HRG stimulation of 184A1 epithelial cells overexpressing HER2** Full KSTAR predictions corresponding to Figure 2A, with phosphoproteomic data obtained from (Wolf-Yadlin et al., Molecular Systems Biology, 2006). Epithelial cells expressing normal HER2 levels (Parental, P) or overexpressing HER2 (24H) were stimulated with EGF or HRG and phosphorylation was measured at 0, 5, 10 and 30 minutes. For each condition, sites with abundance ratios greater than 0.8 were used as evidence, where ratios were relative to the Parental, 5 minute EGF condition (P_EGF_5(min)). Kinase were sorted using hierarchical clustering with Ward linkages. Parental cell stimulated with EGF corresponds to the same culture conditions measured in Figure S3.1

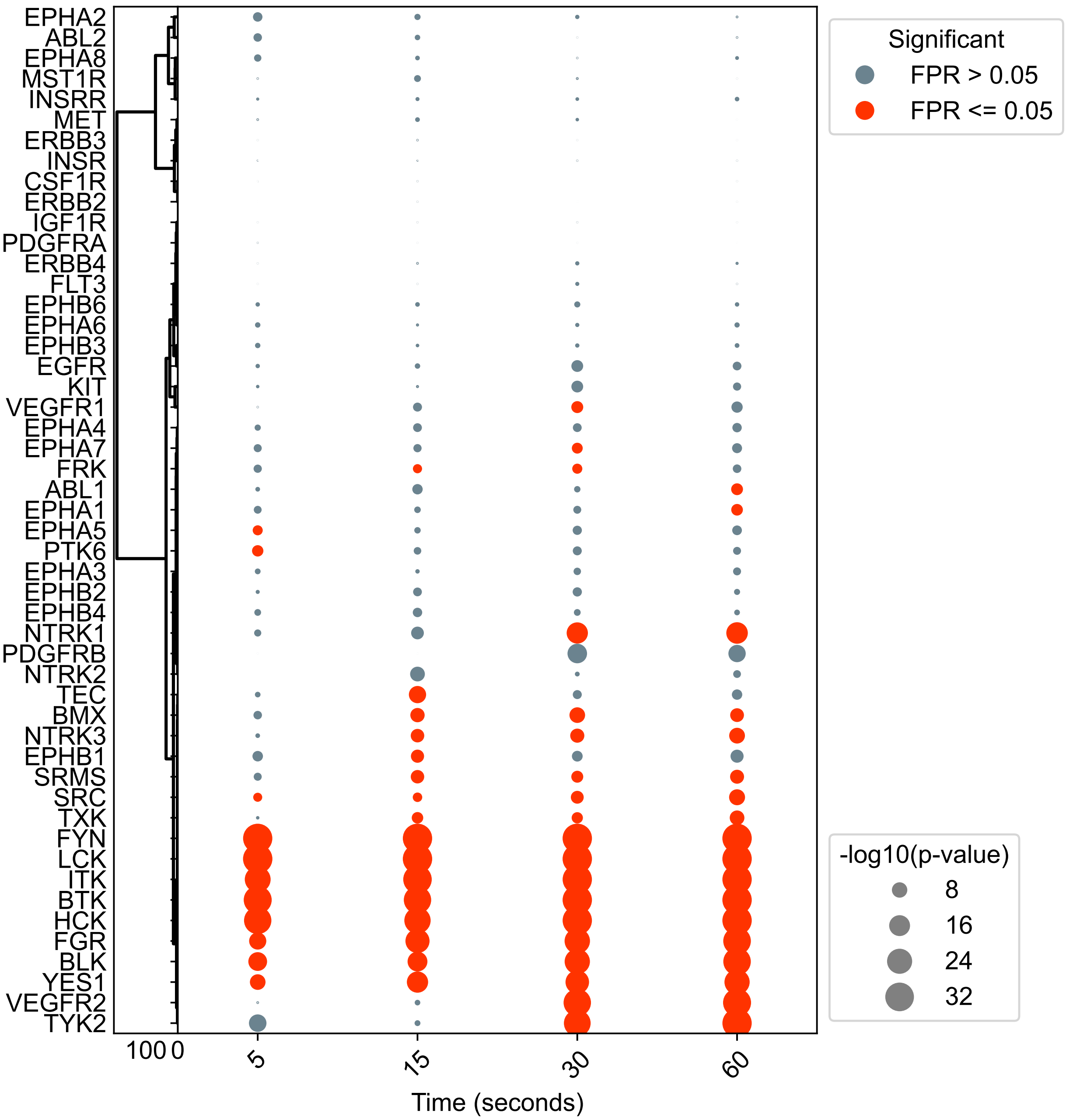


**Figure S3.3. TCR activation of Jurkat cells** Full KSTAR predictions corresponding to Figure 2B based on data from (Chylek et al., PLoS ONE, 2014). Jurkat cells were stimulated to activate T-cell receptor signaling and phosphorylation was measured at 0, 15, 30, and 60 seconds. For each condition, sites with abundance ratios greater than 0.2 were used as evidence, where ratios were relative to the 0 minute timepoint. Kinases were sorted using hierarchical clustering with Ward linkages

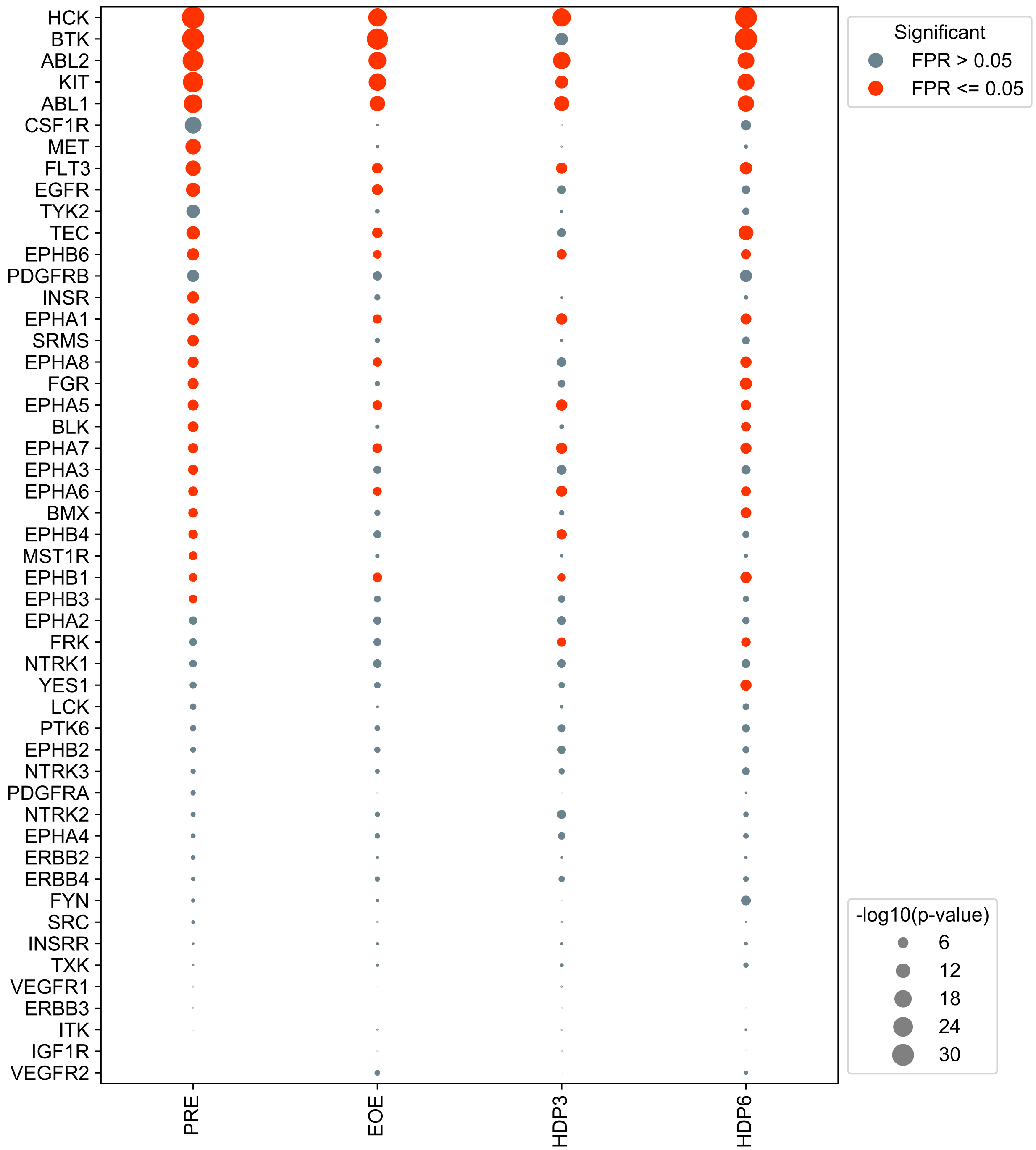


**Figure S3.4. BCR-ABL inhibition by dasatanib** Full KSTAR predicitions corresponding to Figure 2C, based on data from (Asmussen et al., Cancer Discovery, 2013). K562 chronic myeloid leukemia (CML) cell line, which contains the BCR-ABL fusion protein, was treated with dasatanib, an ABL inhibitor, for 20 minutes prior to drug washout. PRE refers to pre-treatment, EOE refers to the end of treatment, HDP3 refers to 3 hours post drug washout, and HDP6 refers to 6 hours post drug washout. For each condition, sites with abundance ratios greater than 0.5 were used as evidence, where ratios were relative to the 0 minute timepoint. Kinases were sorted using hierarchical clustering with Ward linkages.

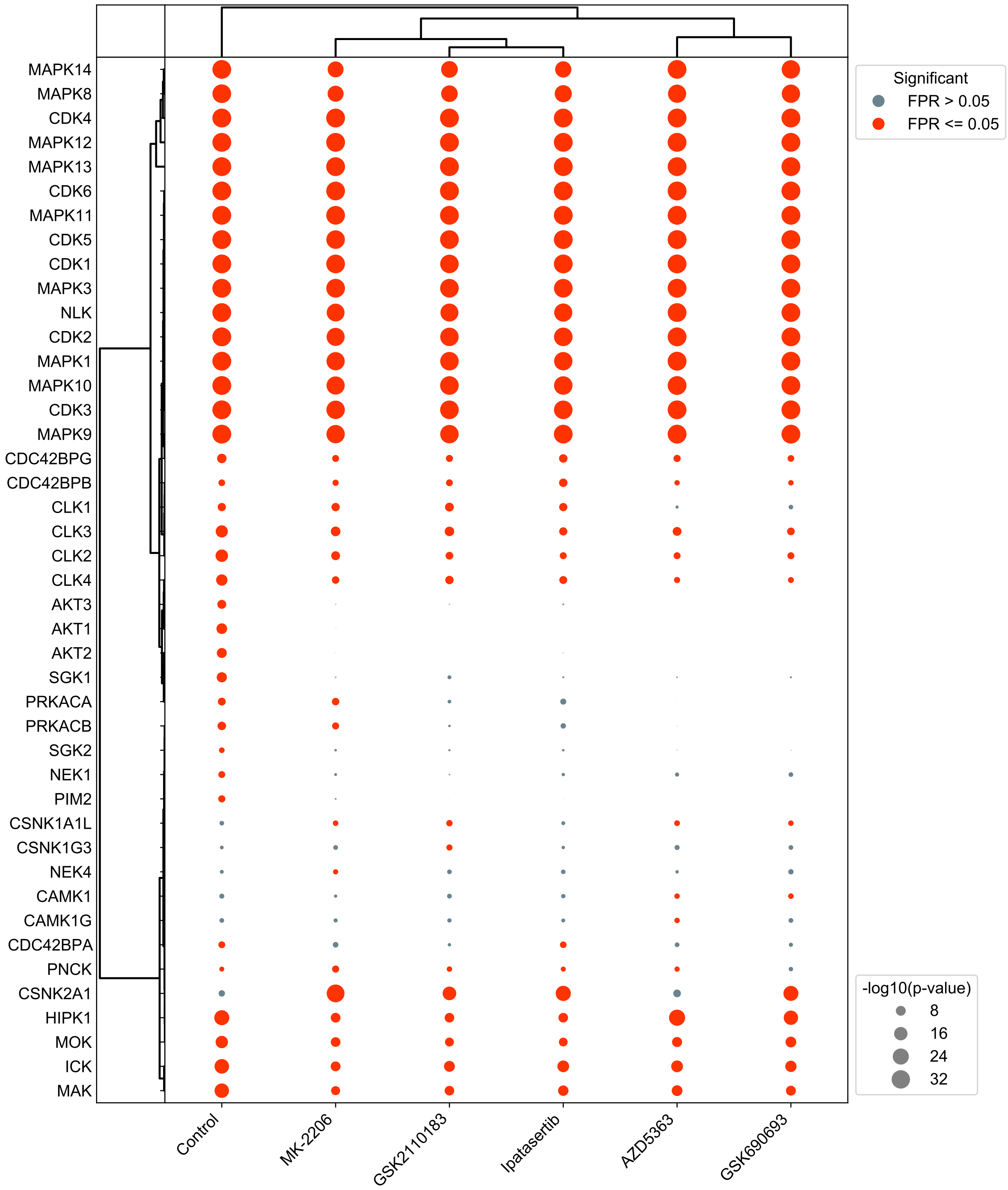


**Figure S3.5. AKT Inhibition with breast cancer cells with 5 different AKT inhibitors**. Full KSTAR predictions corresponding to Figure 2E, based on data from Wiechmann et al. (ACS Chemical Biology, 2021). BT-474 breast cancer cells were treated with one of five different inhibitors (GSK2110183, Ipatasertib, AZD5363, GSK690693, MK-2206). All of the inhibitors work by directly binding the ATP binding pocket except for MK-2206, which is an allosteric inhibitor. For each inhibitor condition, sites with abundance ratios greater than or equal to 1 were used as evidence, relative to pre-treatment contol. Kinases and conditions were sorted using hiearchical clustering with Ward linkages.

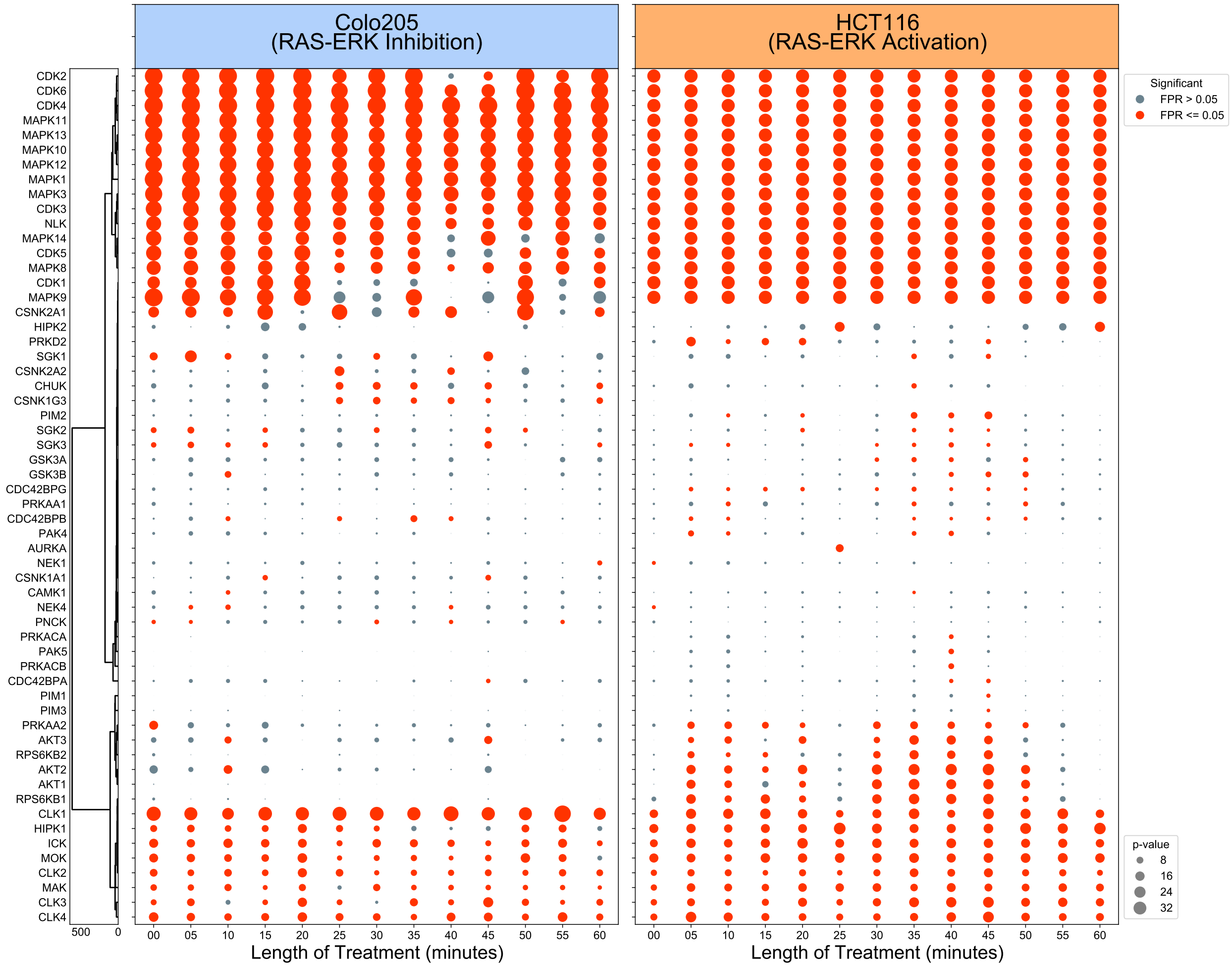


**Figure S3.6. Paradoxical MAPK activation after BRAF inhibition** Full KSTAR predictions corresponding to Figure 2F, based on data from (Kubiniok et al., Molecular and Cellular Proteomics, 2017). Colorectal cancer cell lines Colo205 ($BRAF^{V600E}$ mutation) or HCT116 (KRAS mutation) were treated with the BRAF inhibitor, vemurafenib, over the course of 60 minutes. In cells containing KRAS mutations, BRAF inhibition leads to the activation of the MAP-ERK pathway rather than inhibiting the pathway, as is the case for cells with a $BRAF^{V600E}$ mutation. For each timepoint/cell line, sites with abundance greater than median observed abundance were used as evidence. Kinases were sorted using hierarchical clustering with Ward linkages.

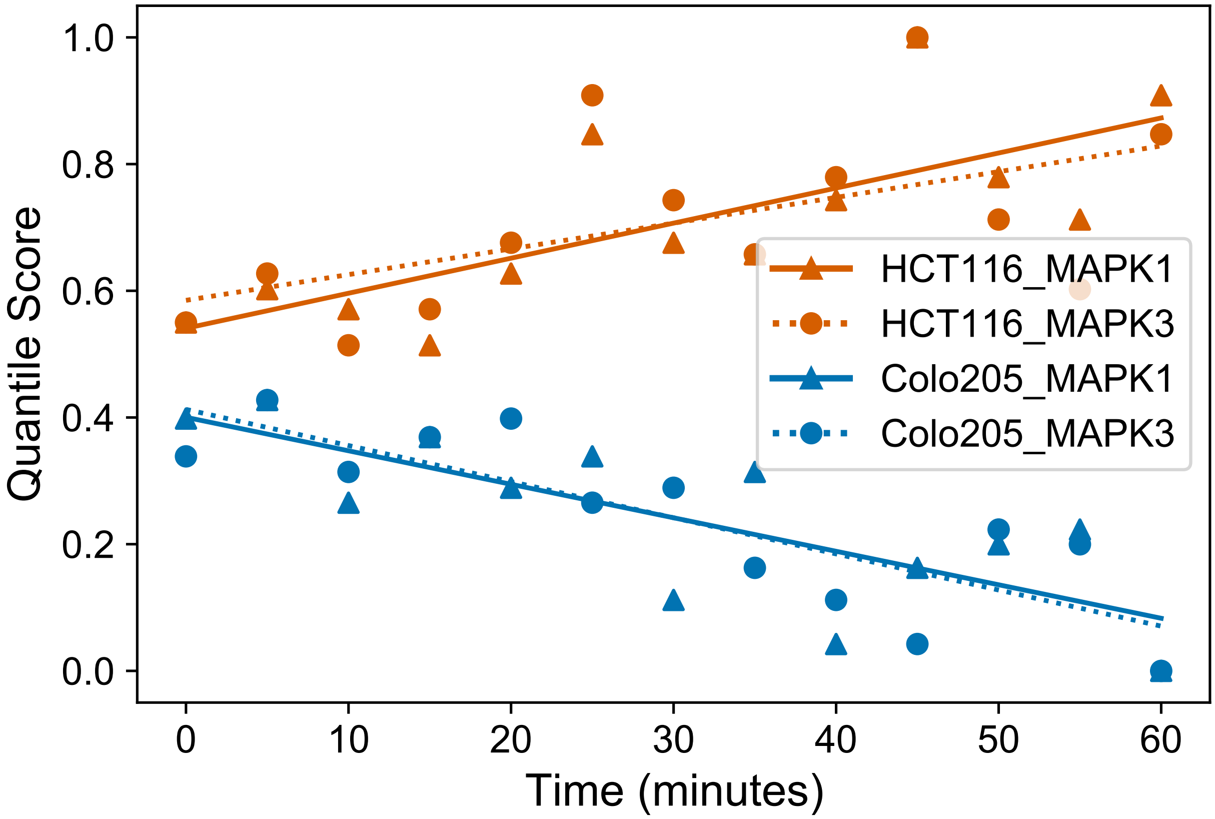


**Figure S3.7. MAPK activity response to BRAF inhibition** Predicted MAPK activity after performing quantile normalization across conditions on the original KSTAR activities corresponding to Figure 2E and Figure S3.6. This demonstrates that despite statistical saturation issues in serine/threonine networks where MAPK/CDK activity is often high, the expected trend is observed (increased MAPK activity in HCT116 cells, decreased MAPK activity in Colo205 cells)
