## Supplemental Figure 4 for "KSTAR: An algorithm to predict patient-specific kinase activities from phosphoproteomic data"

### Supplementary Figures 4: Comparing KSTAR to other available activity inference algorithms

#### Goals

1. Compare the usability and interpretability of various kinase activity inference methods.
2. Compile datasets for use in comprehensive benchmarking of activity inference methods
3. Compare the accuracy of KSTAR to other kinase activity inference methods for both serine/threonine kinases and tyrosine kinases
4. Assess the robustness of algorithms to data loss and the influence of well studied sites on final predictions

#### Methods

As described in the main text, datasets used in the benchmarking analysis were collected from 17 different publications (10 for ST, 7 for Y), described in detail in Table S2. In total, the benchmarking dataset used contained a total of 51 experimental conditions impacting 38 serine/threonine kinases and 19 tyrosine kinases (Figure S4.2). KSTAR, KSEA, PTM-SEA, KARP, and KEA3 were all used to generate predictions about the most enriched/differentially active kinases across the benchmarking dataset (Figure S4.1). Accuracy was calculated based on $P_{hit}$, defined as the fraction of times a kinase expected to perturbed was identified as differentially active, either based on kinase rank or significance. In addition, we looked at the total number of kinases with available predictions for each test condition, per algorithm (Figures S4.3-5).

In addition to accuracy, we also assessed the impact of losing specific sites from the dataset when generating predictions, either by random removal or targeted removal based on the degree of study bias (Figures S4.6 - 9). Starting with 5% loss and continuing to 50% loss (at increments of 5%), the given percent of sites were removed and predictions were regenerated as normal. We then looked at the change in false discovery rate for each prediction, and how this differed between the random and targeted attack. To quantify the differences between these two curves, we defined two metrics: 1) sensitivity to data loss, which is the area under the random attack curve, and 2) sensitivity to study bias, which is the area between the targeted and random attack curve. Given that we are looking at changes to significance of prediction, only KSTAR, KSEA, and PTM-SEA were assessed (KARP and KEA3 do not provide significance of prediction).

#### Table of Contents

**S4.1** - Chart demonstrating how each algorithm can be interpreted in different scenarios, as well as characteristics of each activity score/rank (Page 2)
**S4.2** - Pie charts indicating the distribution of kinases across the benchmarking dataset used here and how the distribution changes when equal weighting of each kinase is applied for $P_{hit}$ calculations. (Page 3)
**S4.3** - Full kinase-specific accuracies for all kinases in the benchmarking dataset (Page 4)
**S4.4** - Number of kinases with predictions for each condition in the benchmarking datasets (Page 5)
**S4.5** - Impact of applying a substrate requirement on the number of kinases with available predictions in KSEA, KARP, and PTM-SEA (Page 6)
**S4.6** - Two individual condition examples of the impact of data loss and study bias on activity predictions, as described in Figure 3 in the main text. (Page 7)
**S4.7** - Average random and targeted loss curves for individual tyrosine kinases (Page 8)
**S4.8** - Average random and targeted loss curves for individual serine/threonine kinases (Page 9)
**S4.9** - Average sensitivity to data loss and study bias for individual kinases (Page 10)

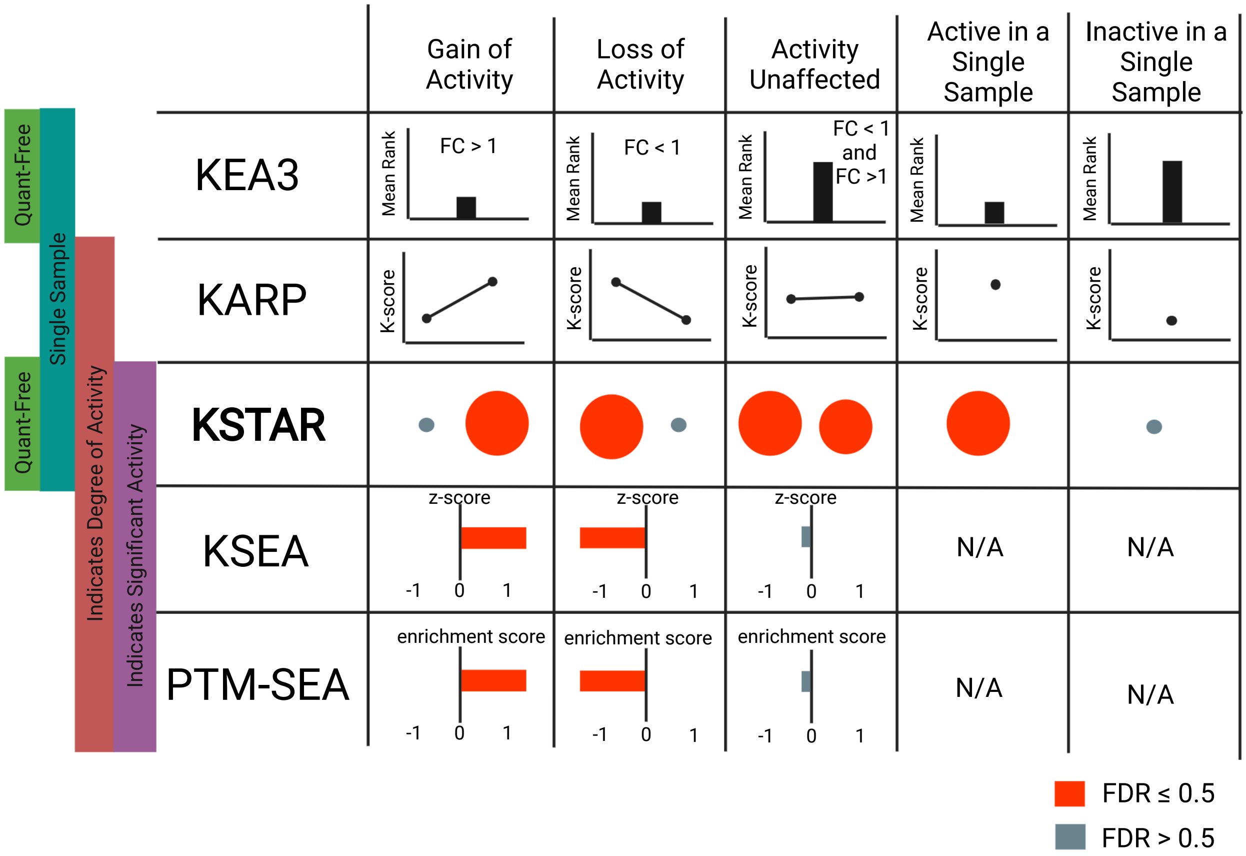


**Figure S4.1. Interpreting kinase activity inference methods.** Kinase activity algorithms differ both in the type of data that can be used and in the way that their output should be interpreted. KSTAR, KARP, and KEA3 are all algorithms that can be used in single sample settings, where differential abundances are not available and an activity score/rank is generated for each sample. While KARP and KSTAR were explicitly designed for use with single sample experiments, KEA3 was largely designed for differential setttings (all three can be used in differential settings as well). On the other hand, KSEA and PTM-SEA require differential abundances. Of the single sample approaches described here, only KSTAR provides both an indicator of the degree of activity and the associated significance of activity. Finally, only KSTAR and KEA3 do not rely on quantification for prediction. Figure created using Biorender.

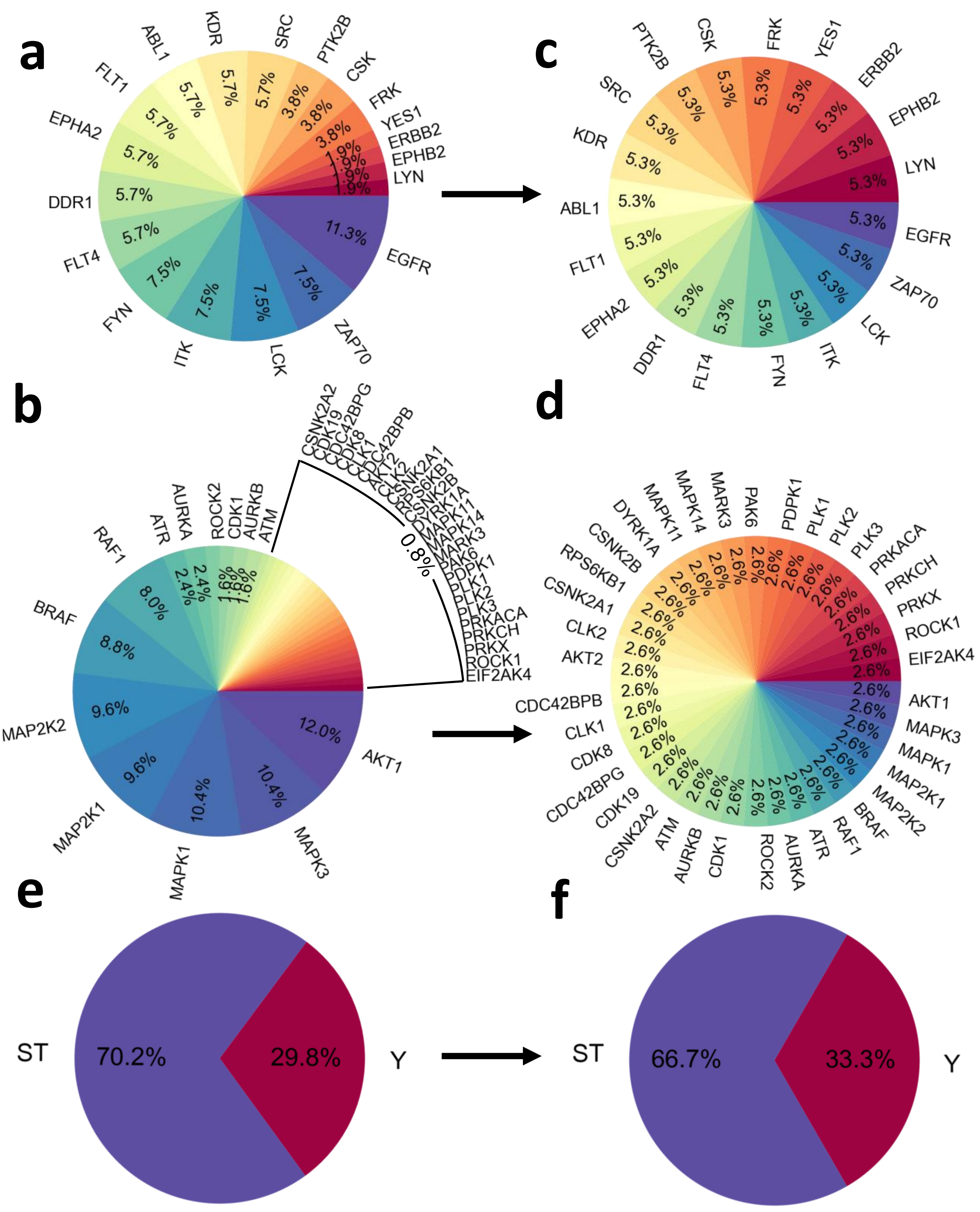


**Figure S4.2. Distribution of kinases in benchmarking dataset. A,B)** To better understand distribution of kinases across the compiled benchmarking dataset, we looked at the percent influence of each kinase, defined by the fraction of conditions for which a kinase is an expected positive. Tyrosine kinases are shown in A, and serine/threonine kinases are shown in B. Certain kinases are overrepresented, such as AKT1 and MAPK1/3 for serine/threonine and EGFR for tyrosine kinases. (caption continued on next page)

**Figure S4.2. C,D)** To avoid certain kinases exhibiting large influence on $P_{hit}$ results, we collapsed each kinase into a single accuracy score ($P_{hit,k}$) and then took the average of each kinase-specific score to obtain the final $P_{hit}$. The impact of this approach can be seen for tyrosine kinases (C) and serine/threonine kinases (D). **E,F)** Distribution of tyrosine and serine/threonine kinases. We found that there was still an overrepresentation of serine/threonine kinases in the dataset both before (E) and after (F) equal weighting was applied. As a result, we separated serine/threonine and tyrosine kinases for benchmarking purposes.

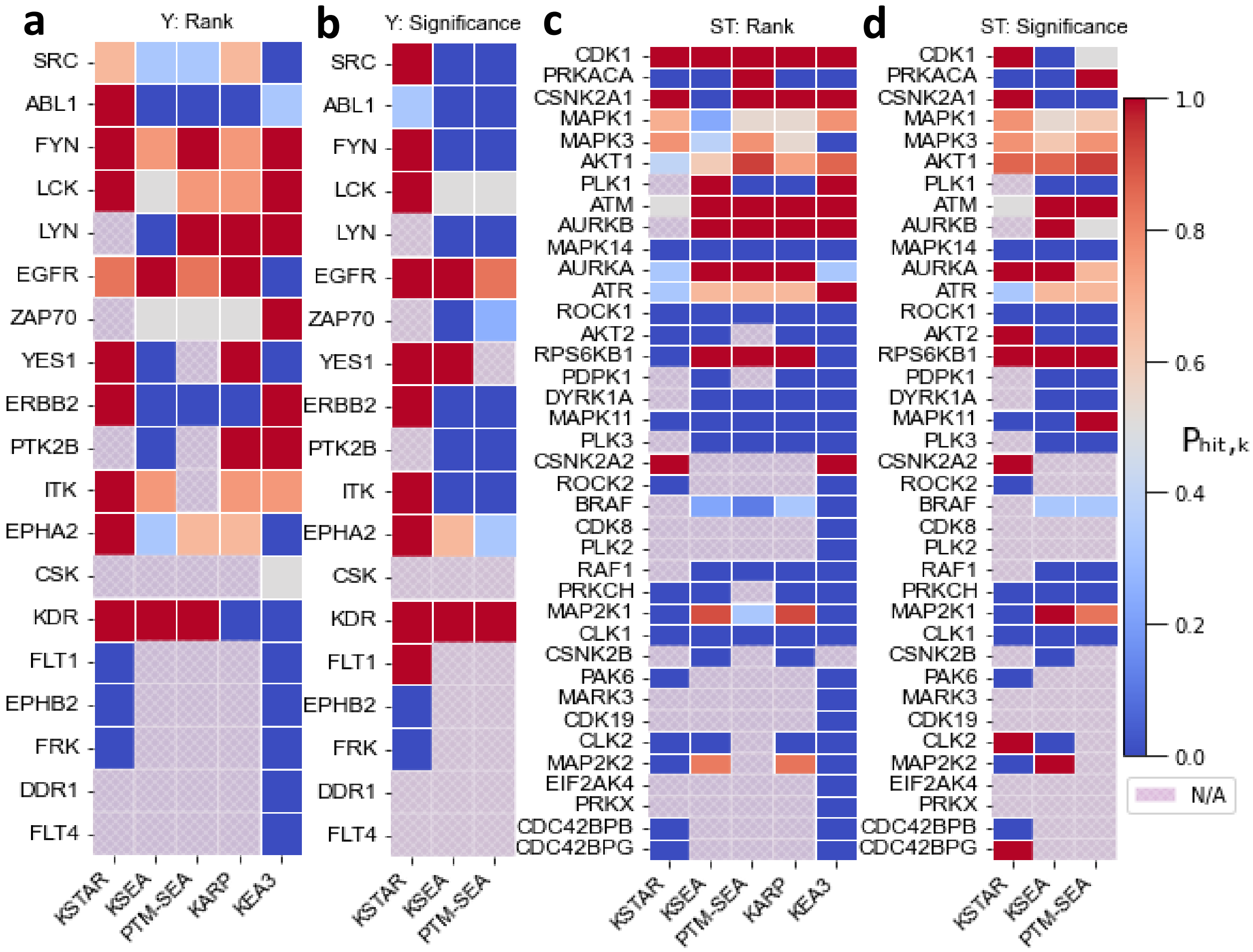


**Figure S4.3. Full results obtained from benchmarking analysis.** Kinase-specific accuracy scores ($P_{hit,k}$) for all kinases perturbed in the benchmarking dataset. All results seen here were used to calculate the final $P_{hit}$ scores in the barplots from Figure 3A in the main text. Kinase predictions were assessed based on rank (found in the top 10 most differentially active kinases) or significance (differential activity has $FDR \leq 0.05$). KARP and KEA3 were only assessed using the rank-based metric. Not all algorithms generated predictions for all kinases, indicated by the light purple squares. **A)** Tyrosine kinase prediction accuracy based on rank. **B)** Tyrosine kinase prediction accuracy based on significance. **C)** Serine/Threonine kinase prediction accuracy based on rank. **D)** Serine/Threonine kinase prediction accuracy based on significance.

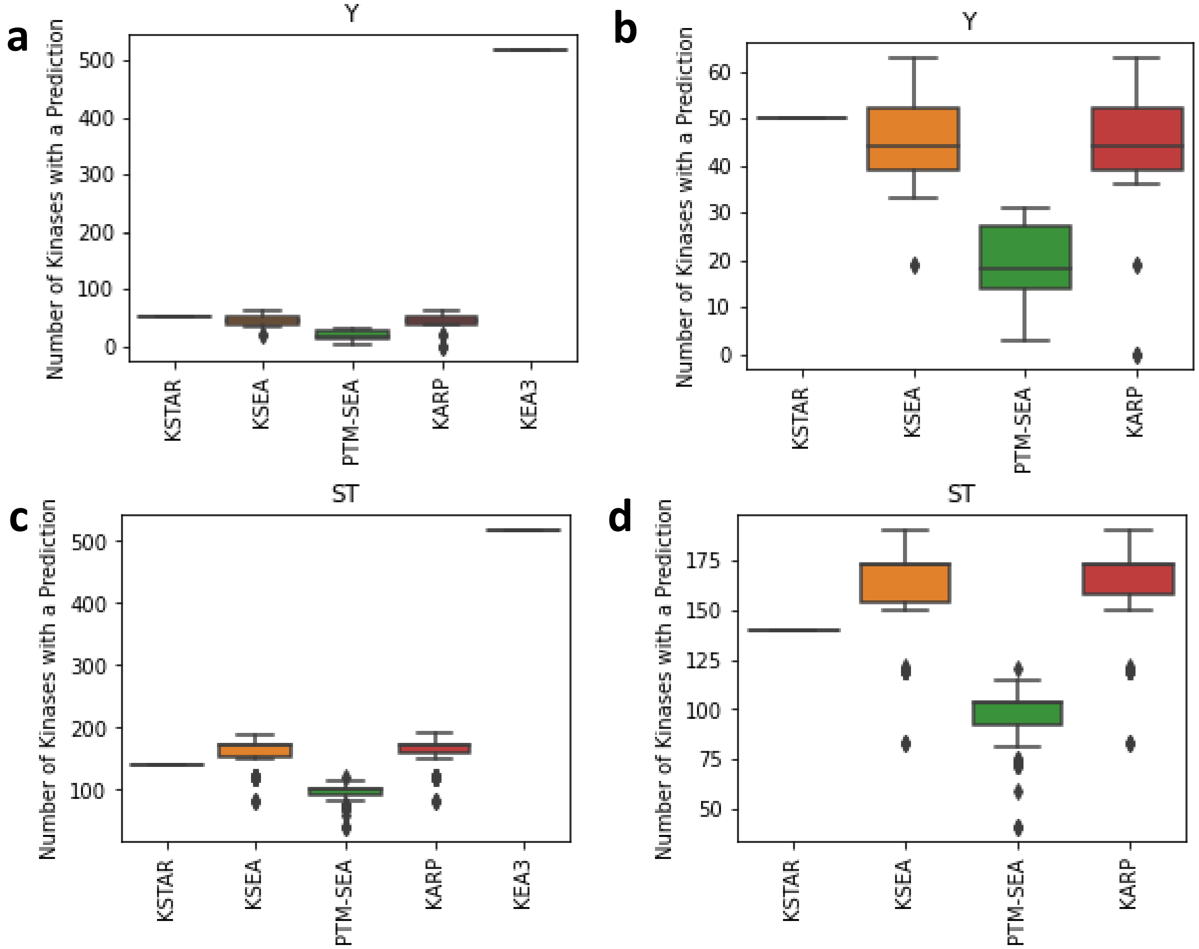


**Figure S4.4. Number of Kinases with Predictions Across Different Activity Inference Algorithms.** Beyond the accuracy of an algorithm, success of kinase activity inference can also be based on the number of different kinases for which predictions are available. Here, we have displayed boxplots indicating the number of different kinases with a prediction in a given dataset (each point corresponds a single dataset result). Because KEA3 generates predictions for most existing kinases, regardless of modification type (tyrosine vs. serine/threonine), we have included plots that omit KEA3. KSTAR generates predictions for 50 tyrosine kinases and 140 serine/threonine kinases for all datasets, while KSEA, PTM-SEA, and KARP depend on the specific phosphorylation sites identified in an experiment, where predictions are only generated when at least one known substrate of a kinase is identified. **A)** Number of kinases with predictions from tyrosine-centric phosphoproteomic datasets, including KEA3. **B)** Number of kinases with predictions from tyrosine-centric phosphoproteomic datasets, omitting KEA3. **C)** Number of kinases with preditions from serine/threonine centric phosphoproteomic datasets, including KEA3. **D)** Number of kinases with prediction from serine/threonine centric phosphoproteomic datasets.

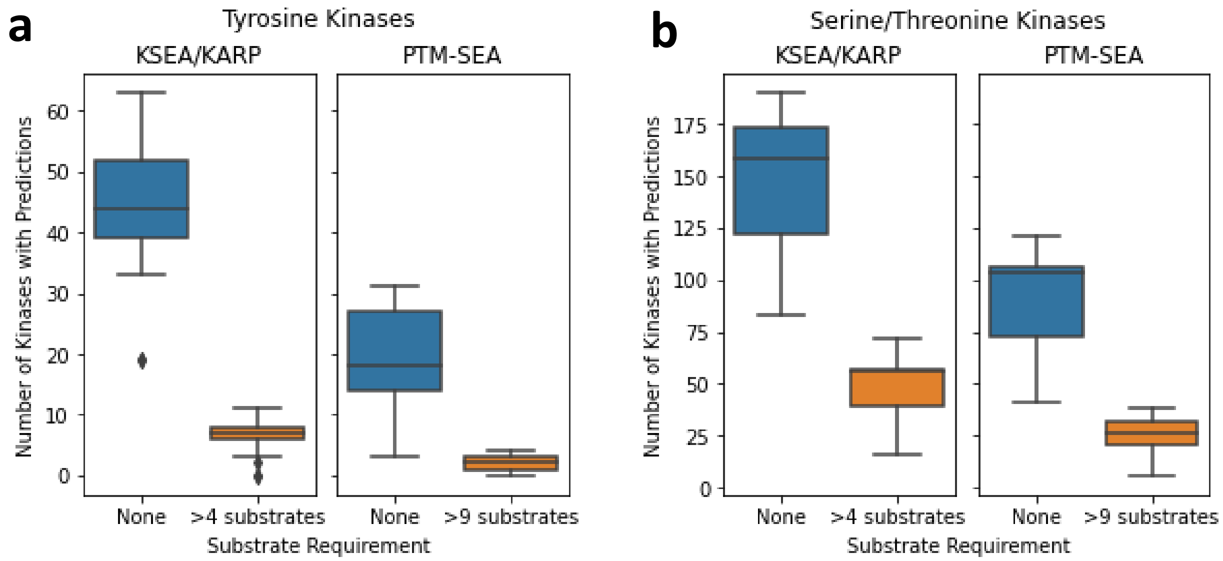


**Figure S4.5. Applying substrate requirements significantly reduces overall kinome coverage of many activity inference algorithms.** For algorithms relying on kinase-substrate annotation, it is common to restrict analyses to kinases with a set number of substrates. For KSEA, which relies on annotation from PhosphoSitePlus, analysis is restricted to kinases with at least five substrates by default. PTM-SEA, which relies on PTMsigDB, restricts analysis to kinases with at least 10 substrates by default. To determine the impact of these requirements, we asked how many kinases had available predictions for each condition in the benchmarking dataset. We found that, in both cases, the substrate requirement significantly reduced the number of available kinase predictions. For tyrosine kinases, there were several cases where less than 5 kinases had available predictions for a given condition. While these predictions are likely to be more robust as more data is associated with each prediction, the lack of kinome coverage significantly reduces the utility of these algorithms. Further, kinases that tend to be connected to well-studied substrates will be more likely to have available predictions, as well studied sites are more likely to be seen in any given experiment (Figure S2). For that reason, all accuracy metrics used in benchmarking were calculated using results without this requirement. **A)** Loss of available kinase predictions for tyrosine kinases when implementing substrate requirements for KSEA (at least five substrates) or PTM-SEA (at least ten substrates). While KARP does not explicitly define a substrate requirement, it also relies on PhosphoSitePlus annotations, so would be equally affected by such a requirement as KSEA. Each data point in the boxplot represents the number of kinases with predictions in a single condition. **B)** Loss of available kinase predictions for serine/threonine kinases when implementing substrate requirements for KSEA (at least five substrates) or PTM-SEA (at least ten substrates). Each data point in the boxplot represents the number of kinases with predictions in a single condition.

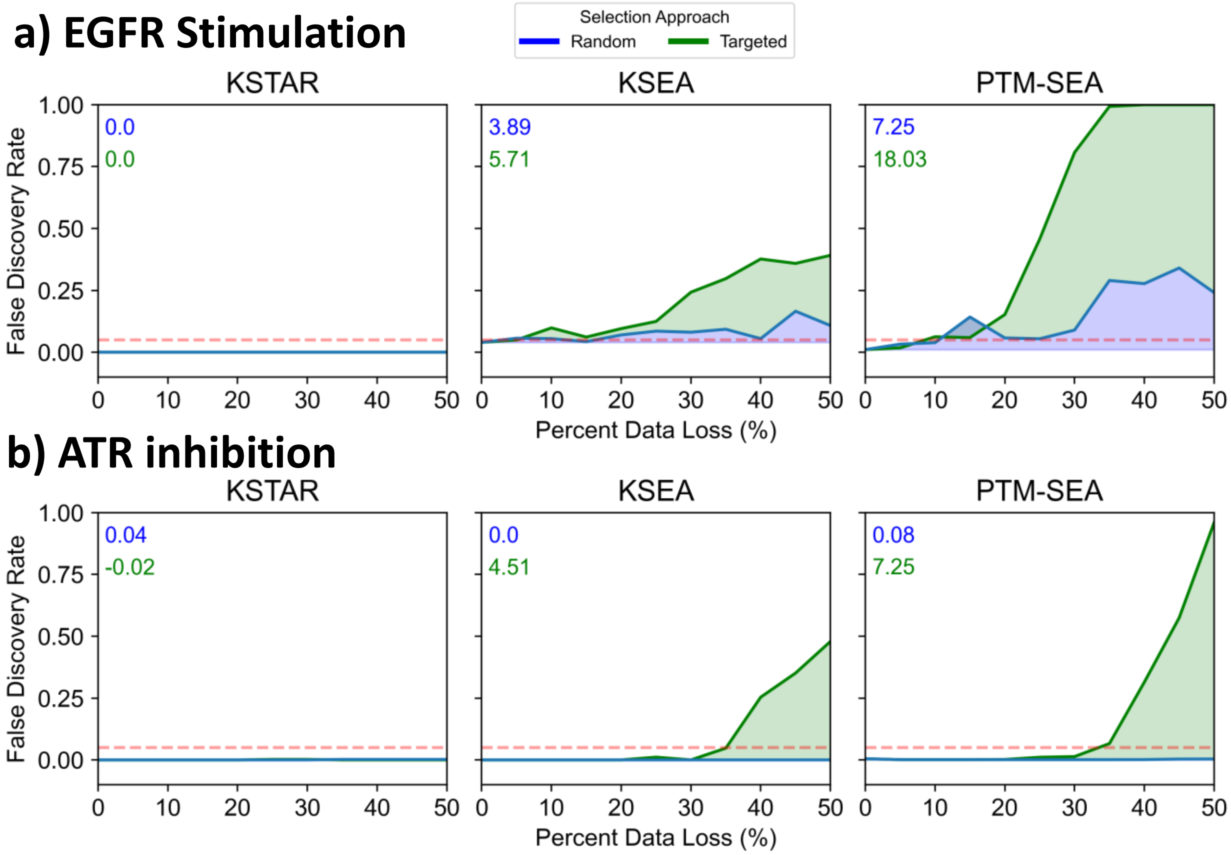


**Figure S4.6. Experiment-specific examples of the influence of data Loss and study bias in individual experiments** Here, we have provided two specific examples of the loss curves obtained for a single condition via either random or targeted attack. As described in the main text, random loss curves are obtained by randomly removing sites (up to 50% of them, done in increments of 5%) and obtaining the average false discovery rate obtained by each algorithm at each data loss amount. Targeted loss curves are obtained in a similar fashion, except that more well studied sites (defined by the number of compendia they are found in) are removed first. A) EGF stimulation condition from Wolf-Yadlin et al. (Proceedings of the National Academy of Sciences, 2007), which is experiment #4 from Table S2, focused on predicted EGFR activity increase after EGF stimulation. **B)** ATR inhibition condition from Beli et al. (Molecular Cell, 2012), which is experiment #51 from Table S2, focused on predicted ATR activity decrease.

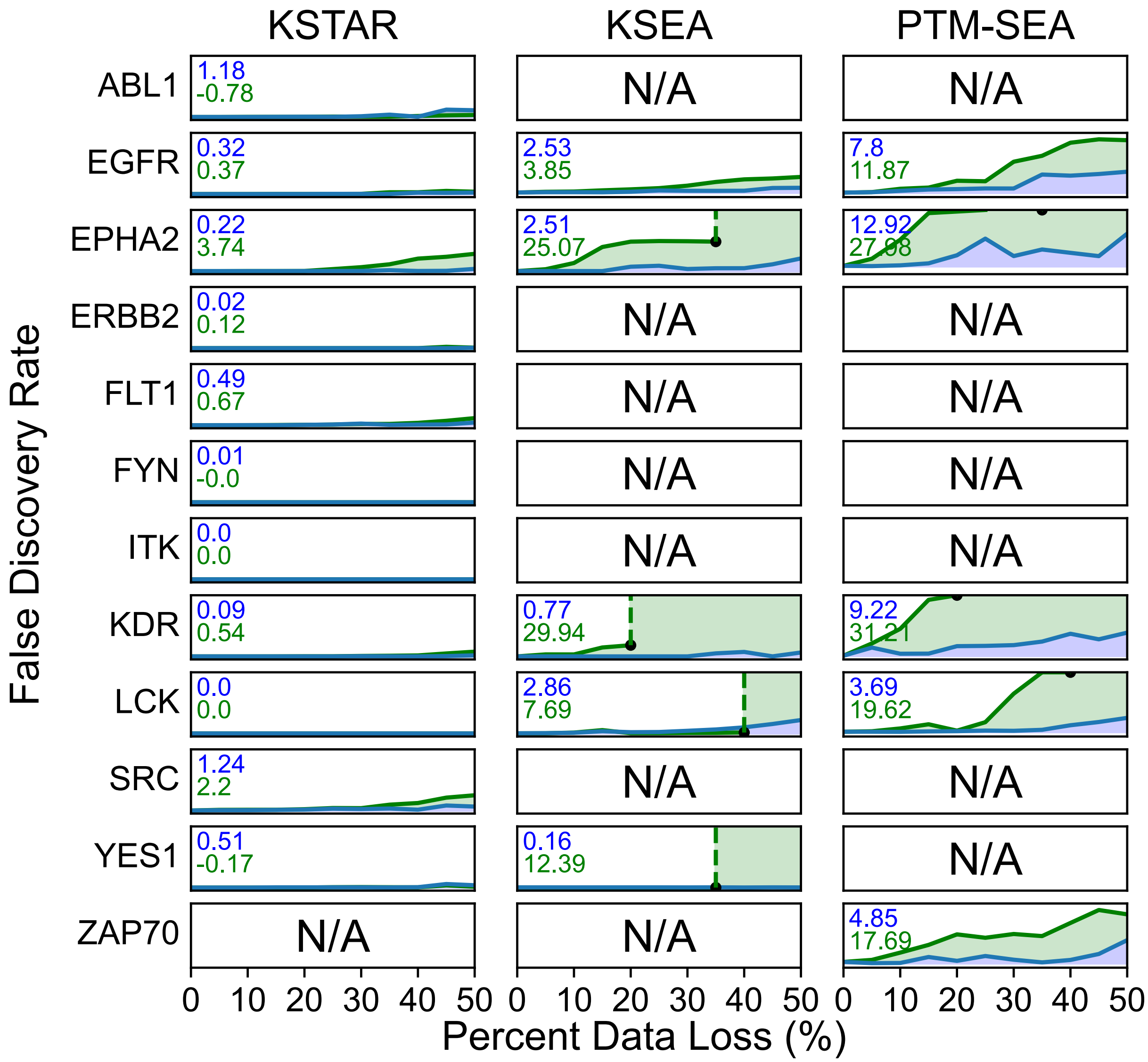


**Figure S4.7. Tyrosine kinase predictions lose significance with increasing data loss.** The average random and targeted loss curves for each tyrosine kinase tested in the data loss experiment (corresponds to Figure 4 of the main text). Rows correspond to a specific kinase, columns correspond to a specific algorithm. The x-axis indicates increasing data loss from 0 to 50%, and the y-axis indicates the false discovery rate of the predicted activity for the kinase of interest, ranging from 0 to 1. The average sensitivity to data loss (blue) and study bias (green) are displayed in the upper left of each plot (these values correspond to heatmap in Figure S4.9). In cases where the curve stops before reaching 50% data loss (indicated by a black point at the stop point), this occurs because there is no longer any available substrates in the data that can be used for prediction in either PhosphoSitePlus (for KSEA) or PTMsigDB (for PTM-SEA).

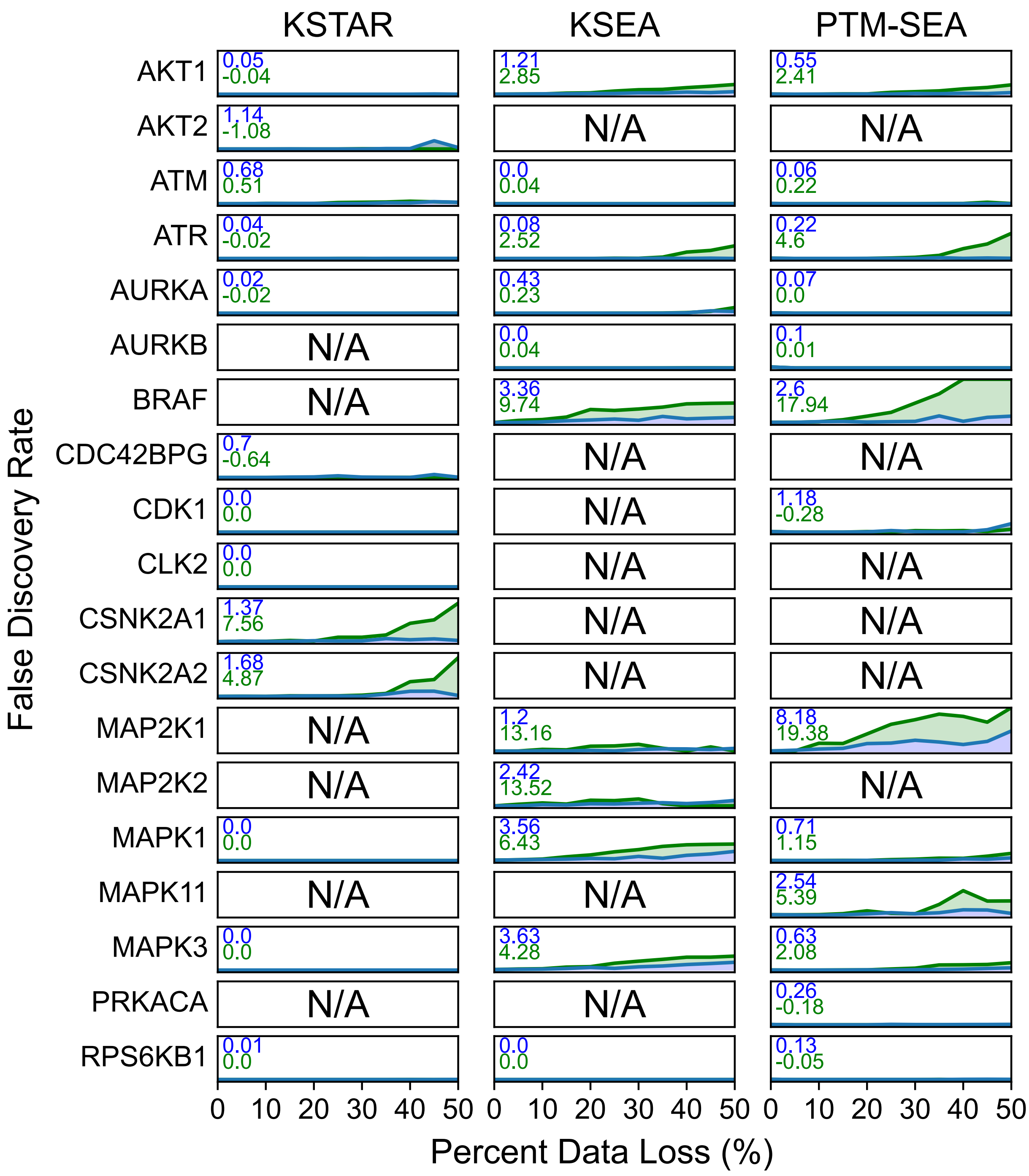


**Figure S4.8. Serine/threonine kinase predictions lose significance with increasing data loss.** The average random or targeted loss curves for each serine/threonine kinase tested in the data loss experiment (corresponds to Figure 4 of the main text). Rows correspond to a specific kinase, columns correspond to a specific algorithm. The x-axis of each plot indicates increasing data loss from 0-50%, and the y-axis indicates the false discovery rate of the predicted activity for the kinase of interest, ranging from 0 to 1. The average sensitivity to data loss (blue) and study bias (green) are displayed in the upper left of each plot (these values correspond to heatmap in Figure S4.9). In cases where the curve stops before reaching 50% data loss, this occurs because there is no longer any available substrates in the data that can be used for prediction in either PhosphoSitePlus (for KSEA) or PTMsigDB (for PTM-SEA).

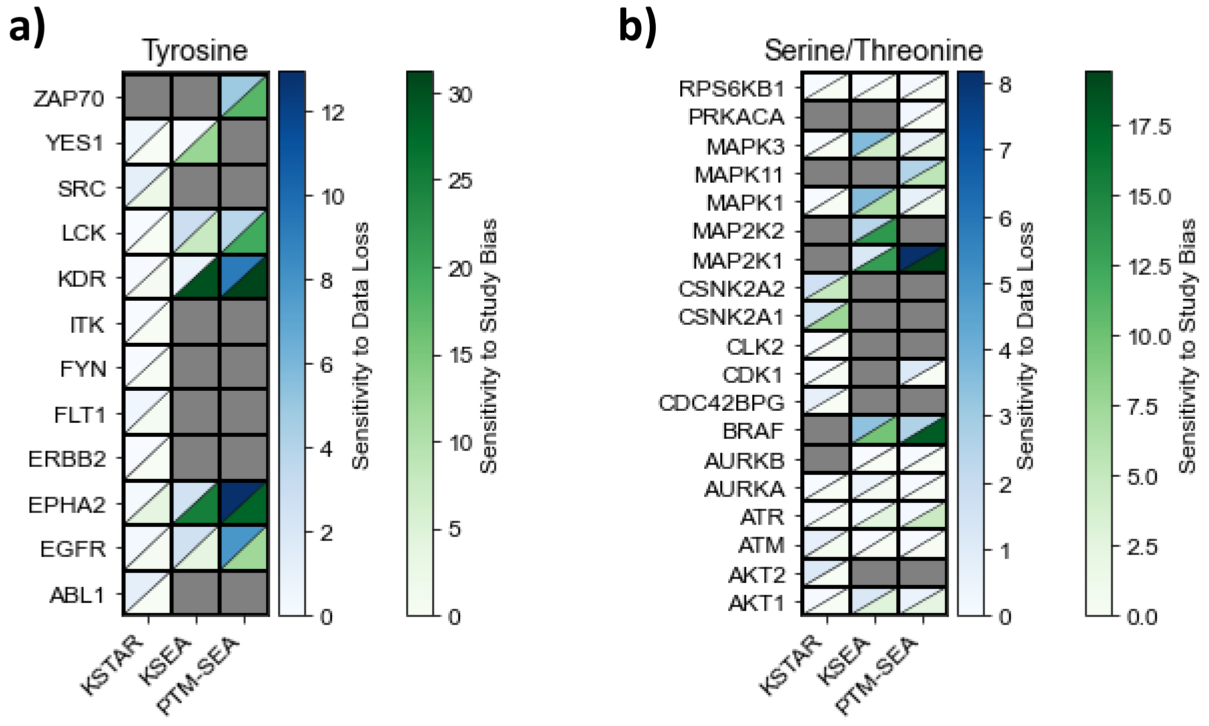


**Figure S4.9. Kinase-specific sensitivity to data loss and study bias for different activity inference methods.** Heatmaps indicating the average sensitivity to data loss and study bias for individual kinases, rather than the whole benchmarking dataset. As described in the main text, "sensitivity to data loss" is defined as the area under the random attack curve, while "sensitivity to study bias" is defined as the difference in the area under the curve between the targeted and random attack. Cells in the heatmap that are gray indicate that predictions were either not available or were not found to be significant in the full dataset. We found that in addition to KSTAR being generally less sensitive to both random and targeted attacks, certain kinase predictions tended to suffer more from targeted attack (removal of well-studied sites), such as KDR (also known as VEGFR2), EPHA2, and MAP2K1. **A)** Tyrosine kinases **B)** Serine/Threonine Kinases
