## Supplemental Figure 5 for "KSTAR: An algorithm to predict patient-specific kinase activities from phosphoproteomic data"

### Supplementary Figures 5: Robustness analysis comparing NSCLC and CML cell lines from independent experiments

#### Goal

Apply KSTAR and KEA3 predictions to multiple different datasets from different labs profiling non small cell lung carcinoma (NSCLC) and/or chronic myeloid leukemia (CML) cell lines to determine if predictions can identify tissue similarities between datasets even when the identified sites differ. These results correspond to Figure 5 in the body of the paper.

#### Methods

A total of 11 datasets from 7 studies and 5 labs were obtained from the original publications. For each dataset, only data corresponding to an untreated cell line was utilized. To generate predictions for each dataset, any site identified in the experiment, regardless of quantification was utilized as evidence. In KSTAR, the phophorylated sites were inputted into the algorithm to produce activity scores and false positive rates. In KEA3, the gene names associated with each phosphorylated site were extracted and inputted into the KEA3 API in python to generate integrated mean ranks, where low mean ranks indicate that the kinase was found to be one of the more enriched kinases in many different protein-protein interaction databases. Spearman rank correlation was used (either with KSTAR activity scores or KEA3 mean ranks) to assess the similarity of predictions across datasets.

#### Table of Contents

**S5.1** Datasets used for the robustness analysis and their characteristics (Page 2)
**S5.2** Full KSTAR activity predictions for each of the 11 datasets and the top ranking kinases in each dataset (Page 3)
**S5.3** Correlation of KEA3 mean ranks across each of the 11 datasets (Page 4)

| Dataset # | Cell Line | Cancer Type | MS Lab | Phosphotyrosine Enriched? | Number of Sites Identified | Reference |
|---|---|---|---|---|---|---|
| 1 | H3255 | NSCLC | Comb | Yes | 466 | A. Guo, et al. Signaling networks assembled by oncogenic EGFR and c-Met. Proceedings of the National Academy of Sciences of the United States of America, 105(2):692–697, 2008. |
| 2 | H3255 | NSCLC | Comb | Yes | 444 | K. Rikova,et al. Global Survey of Phosphotyrosine Signaling Identifies Oncogenic Kinases in Lung Cancer. Cell, 131(6):1190–1203, 2007 |
| 3 | HCC827 | NSCLC | Comb | Yes | 469 | A. Guo, et al. Signaling networks assembled by oncogenic EGFR and c-Met. Proceedings of the National Academy of Sciences of the United States of America, 105(2):692–697, 2008. |
| 4 | HCC827 | NSCLC | Comb | Yes | 452 | K. Rikova, et al. Global Survey of Phosphotyrosine Signaling Identifies Oncogenic Kinases in Lung Cancer. Cell, 131(6):1190–1203, 2007 |
| 5 | H3255 | NSCLC | Comb | Yes | 414 | A. Moritz, et al. Akt - RSK - S6 kinase signaling networks activated by oncogenic receptor tyrosine kinases. Science Signaling, 3(136):1–12, 2010. |
| 6 | HCC827 | NSCLC | Jiminez | Yes | 2586 | R. Beekhof, et al. INKA, an integrative data analysis pipeline for phosphoproteomic inference of active kinases. Molecular systems biology, 15(4):e8250, 2019. |
| 7 | H3255 | NSCLC | Pandey | Yes | 189 | X. Zhang, et al. Quantitative tyrosine phosphoproteomics of Epidermal growth factor receptor (EGFR) tyrosine kinase inhibitor-treated lung adenocarcinoma cells reveals potential novel biomarkers of therapeutic response. Molecular and Cellular Proteomics, 16(5):891–910, 2017. |
| 8 | K562 | CML | Heck | No | 167 | S. Di Palma, et al. Finding the same needles in the haystack? A comparison of phosphotyrosine peptides enriched by immuno-affinity precipitation and metal-based affinity chromatography. Journal of Proteomics, 91:331–337, 2013. |
| 9 | K562 | CML | Shah | Yes | 161 | J. Asmussen, et al. MEK-Dependent Negative Feedback Underlies BCR-ABL-Mediated Oncogene Addiction. Cancer discovery, pages 200–215, dec 2013. |
| 10 | K562 | CML | Jiminez | Yes | 1185 | R. Beekhof, et al. INKA, an integrative data analysis pipeline for phosphoproteomic inference of active kinases. Molecular systems biology, 15(4):e8250, 2019. |
| 11 | K562 | CML | Heck | Yes | 1414 | S. Di Palma, et al. Finding the same needles in the haystack? A comparison of phosphotyrosine peptides enriched by immuno-affinity precipitation and metal-based affinity chromatography. Journal of Proteomics, 91:331–337, 2013. |

**Figure S5.1. Datasets used for robustness analysis** Description of all datasets used for the robustness analysis presented here and in Figure 3. Information includes the specific cell line measured (and corresponding cancer type), the primary mass spectrometry lab that performed the original phosphoproteomic analysis, whether the sample was enriched for phosphotyrosines prior to measurement, and the total number of sites identified in the experiment.

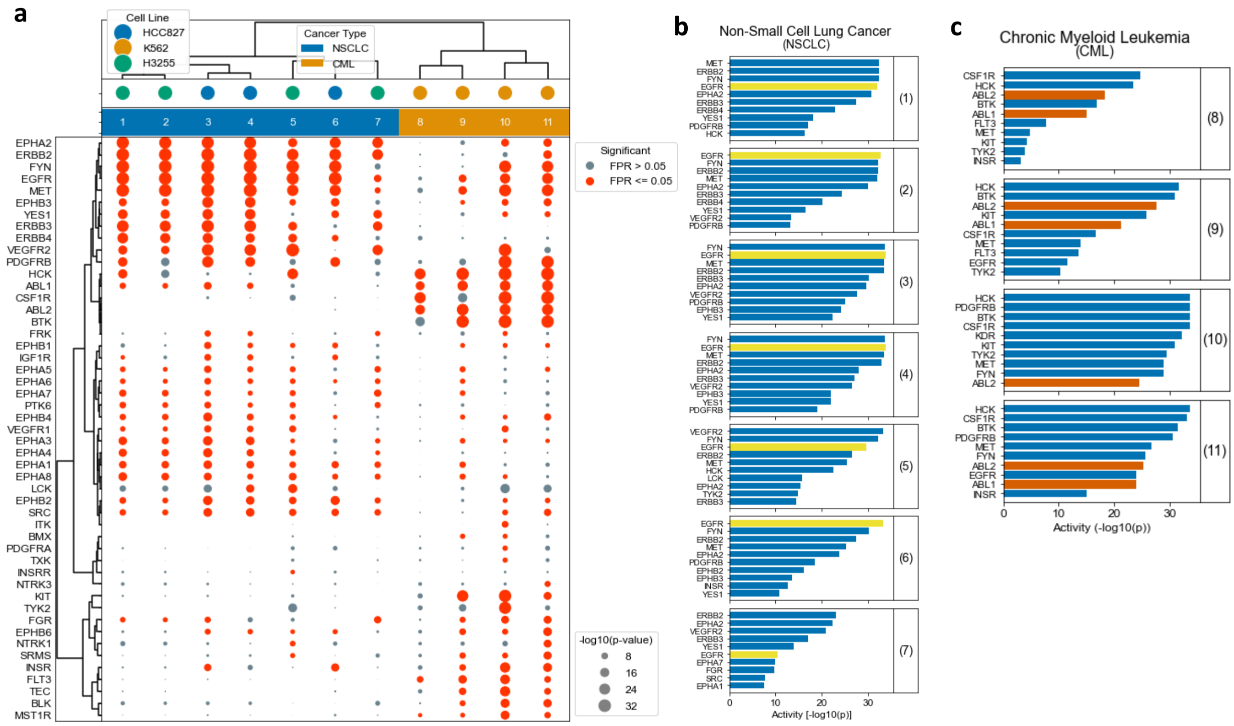


**Figure S5.2. Full KSTAR predictions on NSCLC and CML cell lines** The full KSTAR predictions across all datasets used in robustness analysis. A) Full KSTAR dotplot including activity predictions for all kinases with at least one sample that has significant activity ($FPR <= 0.05$). The size of each dot indicates the degree of predicted activity, with red dots indicating significant activity. Context bars above the plot indicate the cancer type and dataset number. Context dots above the plot indicate the specific cell line measured. Kinases and samples are sorted using hierarchical clustering with ward linkages. B) Top 10 most active kinases in each non small cell lung cancer (NSCLC) dataset, as predicted by KSTAR. Size of the bar indicates the degree of activity. As both H3255 and HCC827 contain activating EGFR mutations, we have the EGFR activity bar in yellow. C) Top 10 most active kinases in each chronic myeloid leukemia (CML) dataset, as predicted by KSTAR. ABL1 and ABL2 are highlighted in red, as the K562 cell lines contains the BCR-ABL fusion protein.

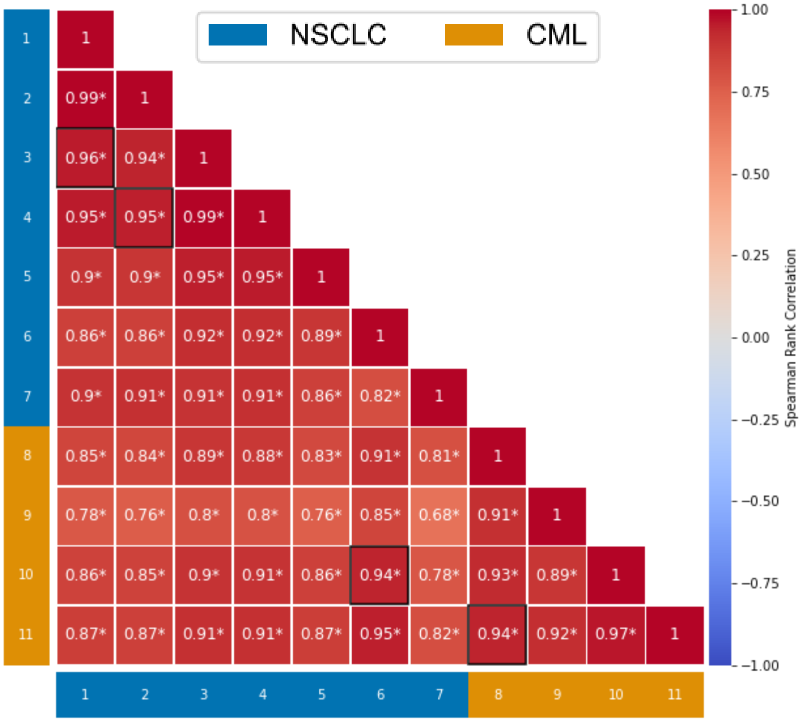


**Figure S5.3. KEA3 predictions on NSCLC and CML cell lines** Heatmap indicating the pairwise correlation between KEA3 rankings obtained from each dataset used in the robustness analysis, based on Spearman correlation coefficient. KEA3 ranks are calculated by reducing the predictions to only include the 50 tyrosine kinases predicted in KSTAR, and then ranked according to KEA3 mean rank. Datasets are sorted according to the same sorting used for the KSTAR heatmap in Figure 3 and in Figure S5.2. Heat cells surrounded by a black box indicate that datasets were obtained from the same study. Overall, this demonstrates that KEA3 was unable to clearly differentiate between the tissues across datasets, with most predictions exhibiting high correlation between each other, regardless of tissue type.
