## Supplemental Figure 6 for "KSTAR: An algorithm to predict patient-specific kinase activities from phosphoproteomic data"

### Supplementary Figures 6: Full analysis results of breast cancer phosphoproteomic datasets

#### Goal

1. Assess the ability of kinase activity inference approaches to predict treatment response in the clinic, particularly in the context of HER2 activity in breast cancer
2. Compare the efficacy of KSTAR, KEA3, and KSEA in the clinical setting

#### Methods

In this supplement, we have provided the KSTAR predictions for all kinase and samples discussed in Figure 6 of the main text, focused on predicting kinase activity in breast tumor biopsies. For the microscaled biopsies of HER2+ patients (Satpathy et al., Nature Communications, 2020), we have included multiple different dotplots to seperate between the pre- and post-treatment data and provide the predictions for serine/threonine kinases in pre-treatment samples (to illustrate that serine/threonine kinase activity profiles do not appear to differentiate between responders and nonresponders).

In addition, where relevant, we have compared our KSTAR results to the results obtained by either KSEA (Figure S6.2) or KEA3 (Figures S6.2, S6.8, S6.10, and S6.11). We found that KSEA results from the tumor biopsies tended to be nonrobust and difficult to apply in this setting (due to low numbers of known substrates and reliance on quantification), so we did not continue use of KSEA after application to the CPTAC dataset (Mertins et al., Nature, 2016). To make KEA3 more directly comparable to KSTAR results, the list of kinases with predictions from KEA3 was reduced to include only kinases that also have predictions in KSTAR. As all predictions are based on the abundance of phosphorylated tyrosines (except in S6.6), this also had the effect of removing serine/threonine kinases from KEA3 predictions, which are not relevant to the actual input data. The KEA3 rankings displayed throughout this supplement are then obtained based on the KEA3 mean ranks, where the 10th ranked kinase had the 10th smallest mean rank across the 50 tyrosine kinases with predictions in KSTAR.

#### Table of Contents

**S6.1** - Full KSTAR predictions on CPTAC phosphoproteomic analysis of 77 Breast cancer patients, corresponding to Figure 4A (Page 2)
**S6.2** - KSTAR, KSEA, and KEA3 predictions in the 107 Breast cancer patients in the CPTAC dataset for ERBB2/HER2 and their correlation with HER2 status. Corresponds to Figure 4A (Page 3)
**S6.3** - Full KSTAR predictions on phosphoproteomic analysis of PDX models of breast cancer, corresponding to Figure 4B (Page 4)
**S6.4** Full KSTAR tyrosine kinase activity predictions for microscaled biopsies of HER2+ patients, corresponding to Figure 4C (Page 5)
**S6.5** Only pre-treatment KSTAR tyrosine kinase activity predicitons for microscaled biopsies of HER2+ patients, corresponding to Figure 4C (Page 6)
**S6.6** Only pre-treatement KSTAR serine/threonine kinase activity predictions for microscaled biopsies of HER2+ patients, corresponding Figure 4C (Page 7)
**S6.7** - Patient-specific KSTAR activity predictions for ERBB2 and EGFR in non-pathologically complete responders (non-PCR) from microscaled biopsies, corresponding to Figure 4C (Page 8)
**S6.8** - Patient-specific KEA3 rankings for ERBB2 and EGFR in non-pathologically complete responders (non-PCR) from microscaled biopsies, corresponding to Figure 4C (Page 9)
**S6.9** - Patient-specific KSTAR activity predictions for ERBB2 and EGFR in pathologically complete responders (PCR) from microscaled biopsies, corresponding to Figure 4C (Page 10)
**S6.10** - Patient-specific KEA3 rankings for ERBB2 and EGFR in pathologically complete responders (PCR) from microscaled biopsies, corresponding to Figure 4C (Page 11)
**S6.11** - Ranking of ERBB2 in microscaled biopsies and its relation to HER2 status, with rankings calculated using either KSTAR activity scores or KEA3. Corresponds to Figure 4C (Page 12)

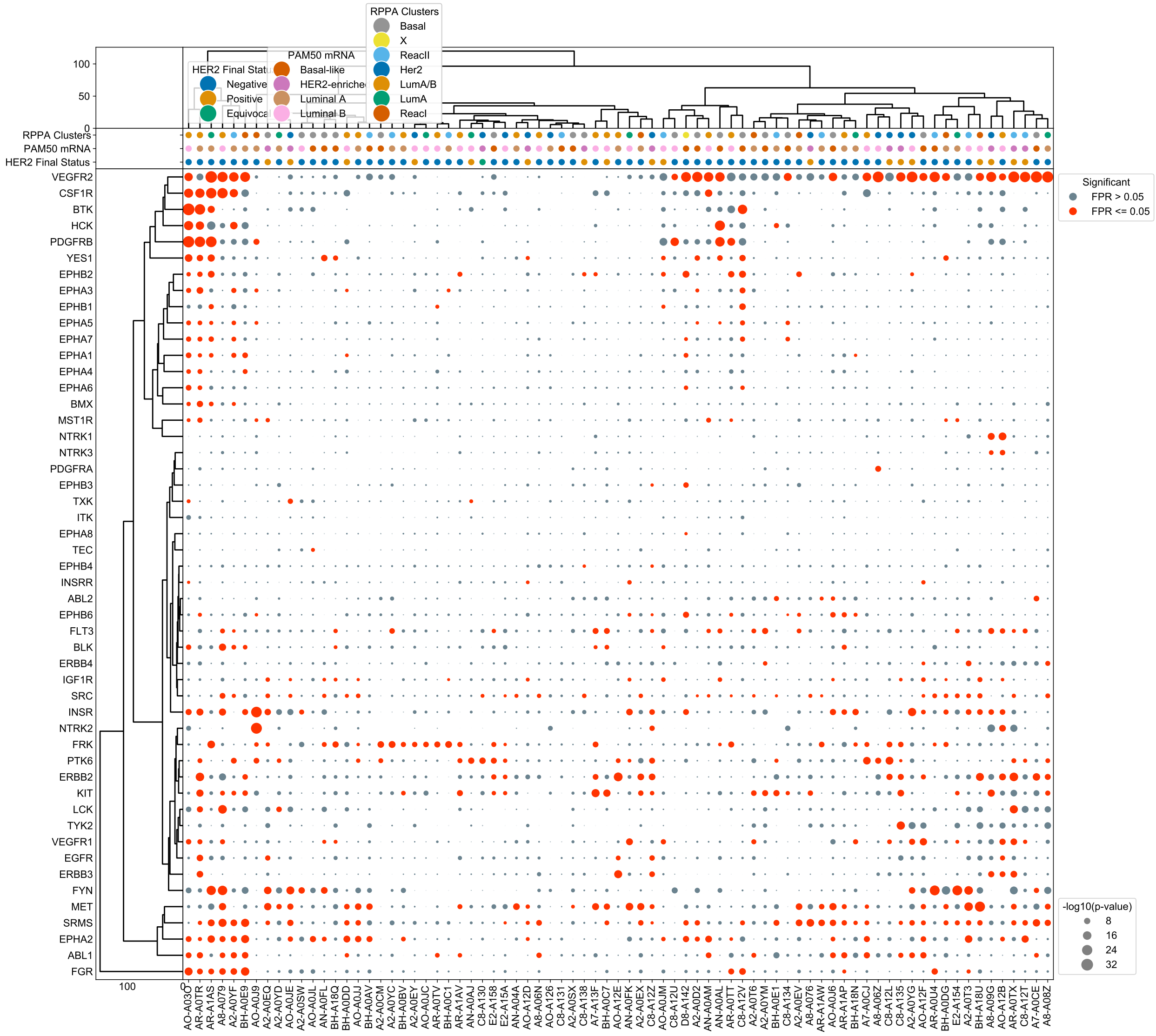


**Figure S6.1. Full KSTAR predictions on CPTAC phosphoproteomic analysis of 77 Breast cancer patients** Dotplot containing all kinase activity predictions for the CPTAC phosphoproteomic dataset (Mertins et al., Nature, 2016), corresponding to Figure 4A. Dot size corresponds the activity score (-log10(p)), and dots are colored based on if the observed activity score has a false positive rate below 0.05. Kinases and samples were sorted using hierarchical clustering with ward linkage. Important clinical attributes of each patient can be found above the main dotplot, including the HER2 status and classification based on either mRNA or protein expression data.

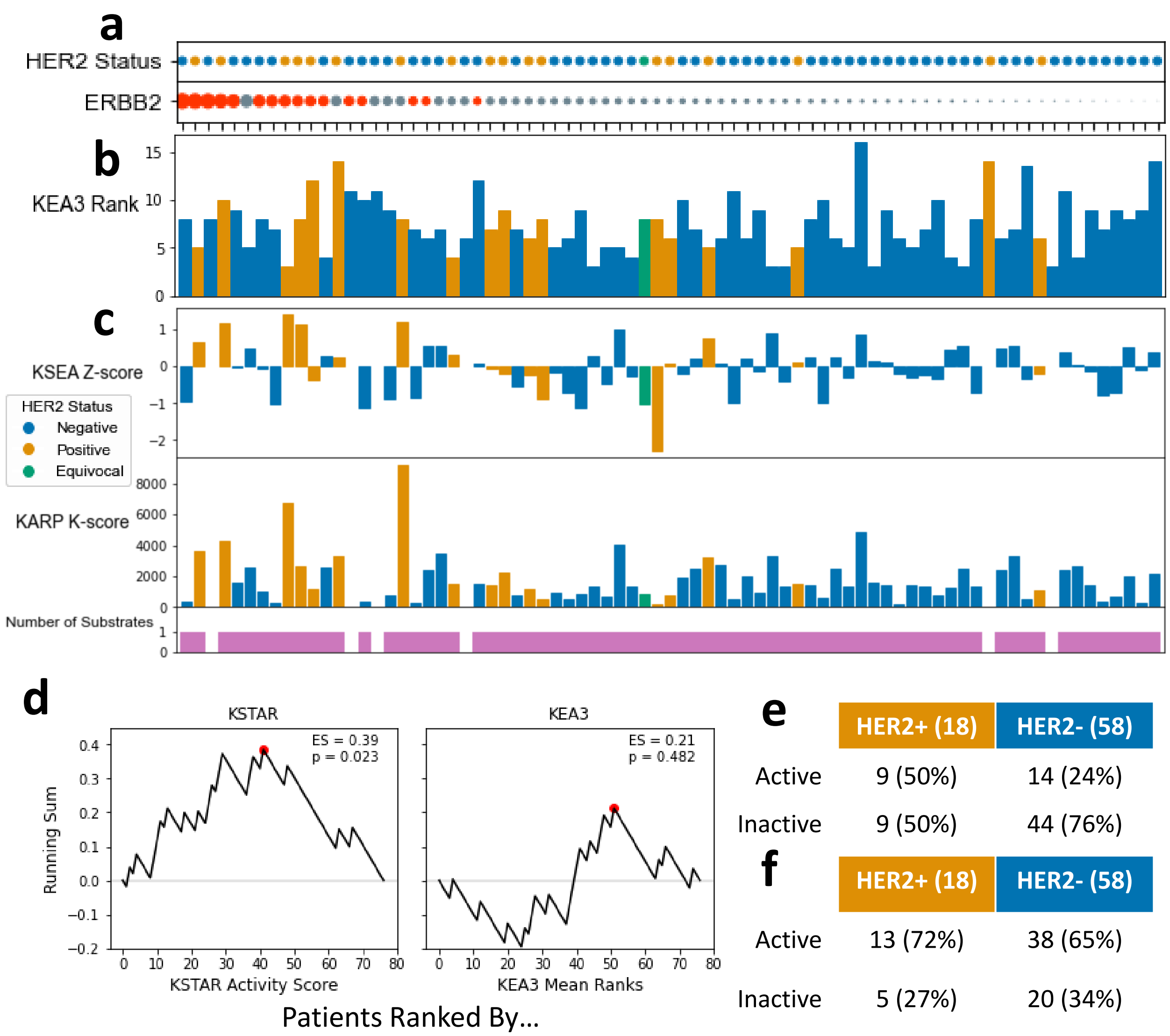


**Figure S6.2. Comparing Kinase Activity Inference Approaches Relationship with HER2 Status** Predicted HER2 activity using different kinase activity inference algorithms (KSTAR, KSEA, KEA3), with patient samples sorted based on KSTAR predicted activity. HER2 status is indicated using the color of the bars/dots depending on plot type. A) KSTAR predictions of ERBB2/HER2 activity, where each where dotsize indicates degree of activity and red samples are significant ($FPR <= 0.05$). This plot is identical to Figure 4A. B) KEA3 ERBB2/HER2 ranking, relative to the 50 kinases that also have predictions in KSTAR. So, for a sample where the rank is 10, ERBB2/HER2 is the 10th most enriched kinase out of 50 other tyrosine kinases. C) KSEA z-scores and number of identified ERBB2 substrates for each CPTAC patient sample. In patients with at least one ERBB2/HER2 substrate, z-score enrichment was calculated with measured log2 transformed abundances. Positive z-scores indicate high ERBB2 activity and negative z-scores indicate low ERBB2 activity, relative to a pooled sample. Empty bars correspond to patients with no identified substrates, as indicated by PhosphoSitePlus. Only 44.2% of patients had KSEA predictions, all of which used only a single site as evidence, LDHA Y10. Significant z-scores are indicated by '*' ($FDR <= 0.05$). D) GSEA-style running sum to test if KSTAR HER2/ERBB2 activity or KEA3 HER2/ERBB2 mean ranks significantly correlates with HER2 status. To calculate the running sum, samples were sorted according to predicted ERBB2 activity obtained from KSTAR or KEA3, with the most ERBB2 active samples at the top of the list. We then moved through the list, adding to the running sum if the sample was HER2+ and subtracting from the running sum if the sample was HER2-. The final enrichment score is equal to the maximum deviation from 0 obtained in the running sum. Statistical significance was then calculated using a null distribution consisting of ES scores obtained when the sample list was sorted randomly. Only KSTAR ERBB2 activity predictions were found to be significantly correlated with HER2 status. E) HER2 status predictions for KSTAR, with active ERBB2/HER2 defined by $score <= 1e-3$. Same table as in Figure 4A. F) HER2 status predictions for KEA3, defining active ERBB2/HER2 by an ERBB2 rank of 8 or higher. This cutoff was determined by identifying the rank cutoff with the best overall F1 score for KEA3.

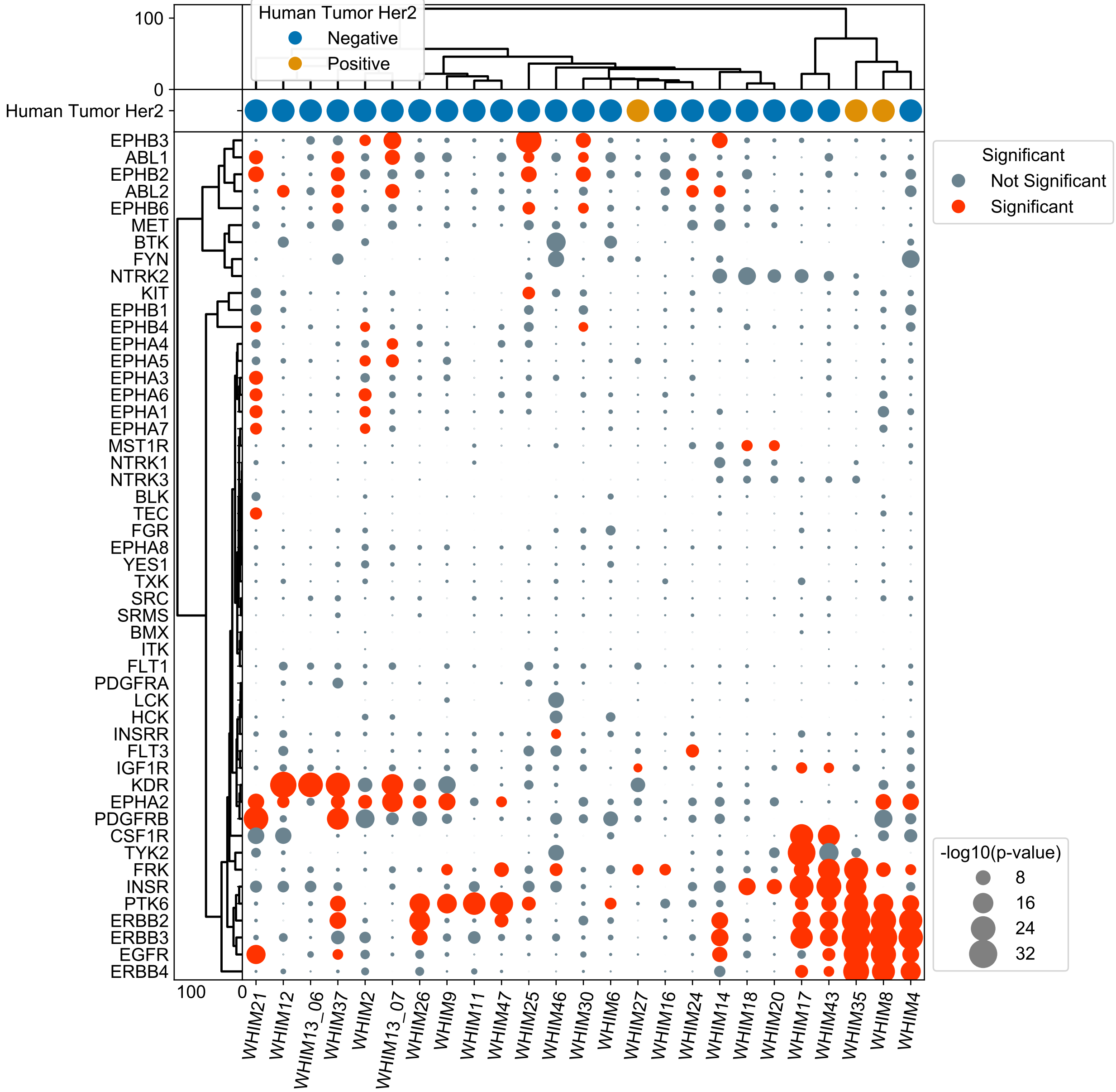


**Figure S6.3. Full KSTAR predictions on PDX Dataset** Dotplot containing all tyrosine kinase activity predictions for the PDX dataset (Huang et al., Nature Communications, 2017), corresponding to Figure 4B. Dot size corresponds the activity score (-log10(p)), and dots are colored based on if the observed activity score has a false positive rate below 0.05. Kinases and samples were sorted using hierarchical clustering with ward linkage. HER2 status of each patient is indicated above the dotplot. 3 of the 25 PDX samples were HER2+, the rest of the samples were HER2-

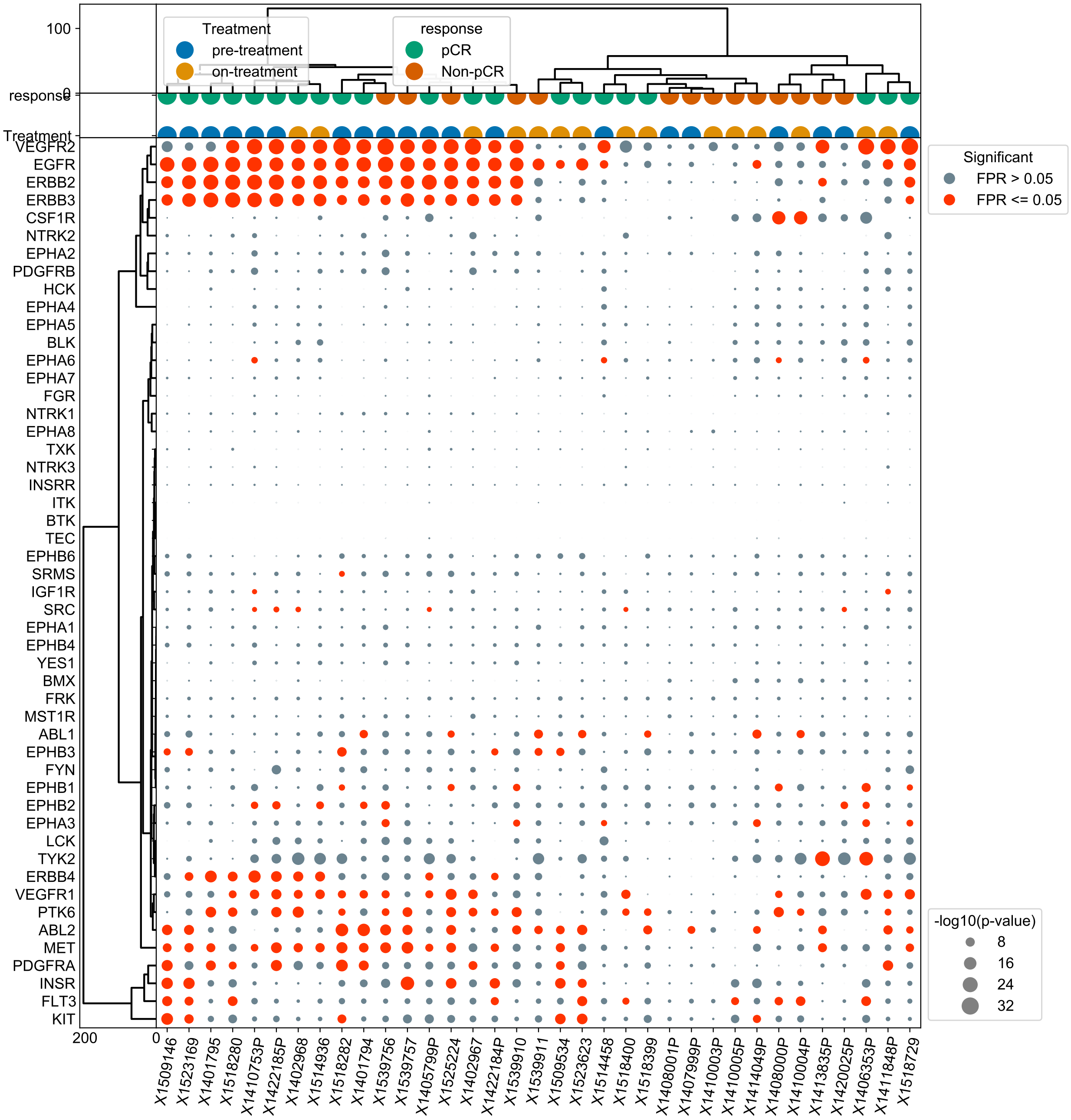


**Figure S6.4. Full KSTAR tyrosine kinase activity predictions on microscaled biopsies** Full dotplot containing all tyrosine kinase activity predictions on Microscaled Biopsies of HER2+ patients for both pre- and post-treatment (Satpathy, Nature Communications, 2020). Corresponds to Figure 4C. Patients were treated with a combination of chemotherapy and HER2 targeted therapy, and post-treatment biopsies were taken 48-72 hours following the beginning of treatment. Dot size corresponds to the activity score (-log10(p)), and dots are colored based on if the observed activity score has a false positive rate below 0.05. Kinases and samples were sorted using hierarchical clustering with ward linkage. Treatment status (pre- vs. post-treatment) and treatment response (responder vs. non-responder) of each sample are indicated with the context dots above the dotplot.

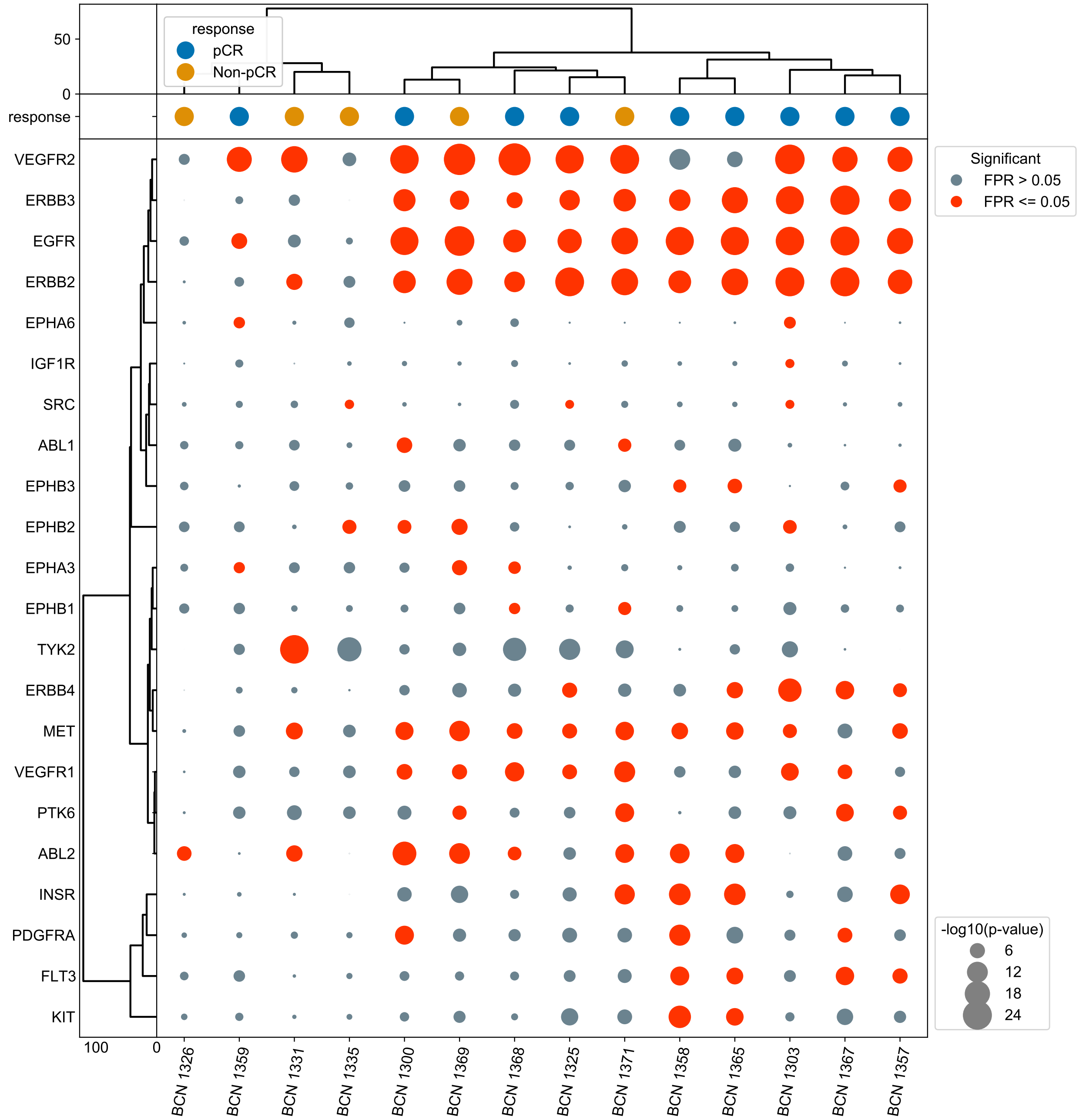


**Figure S6.5. KSTAR tyrosine kinase activity predictions on microscaled biopsies prior to treatment** Full dotplot containing all tyrosine kinase activity predictions on microscaled biopsies of HER2+ patients for only pre-treatment. Corresponds to Figure 4C. Dot size corresponds to the activity score (-log10(p)), and dots are colored based on if the observed activity score has a false positive rate below 0.05. Kinases and samples were sorted using hierarchical clustering with ward linkage. Treatment response to combination therapy of chemotherapy and anti-HER2 therapy (responder/pCR vs. non-responder/non-pCR) of each sample are indicated with the context dots above the dotplot.

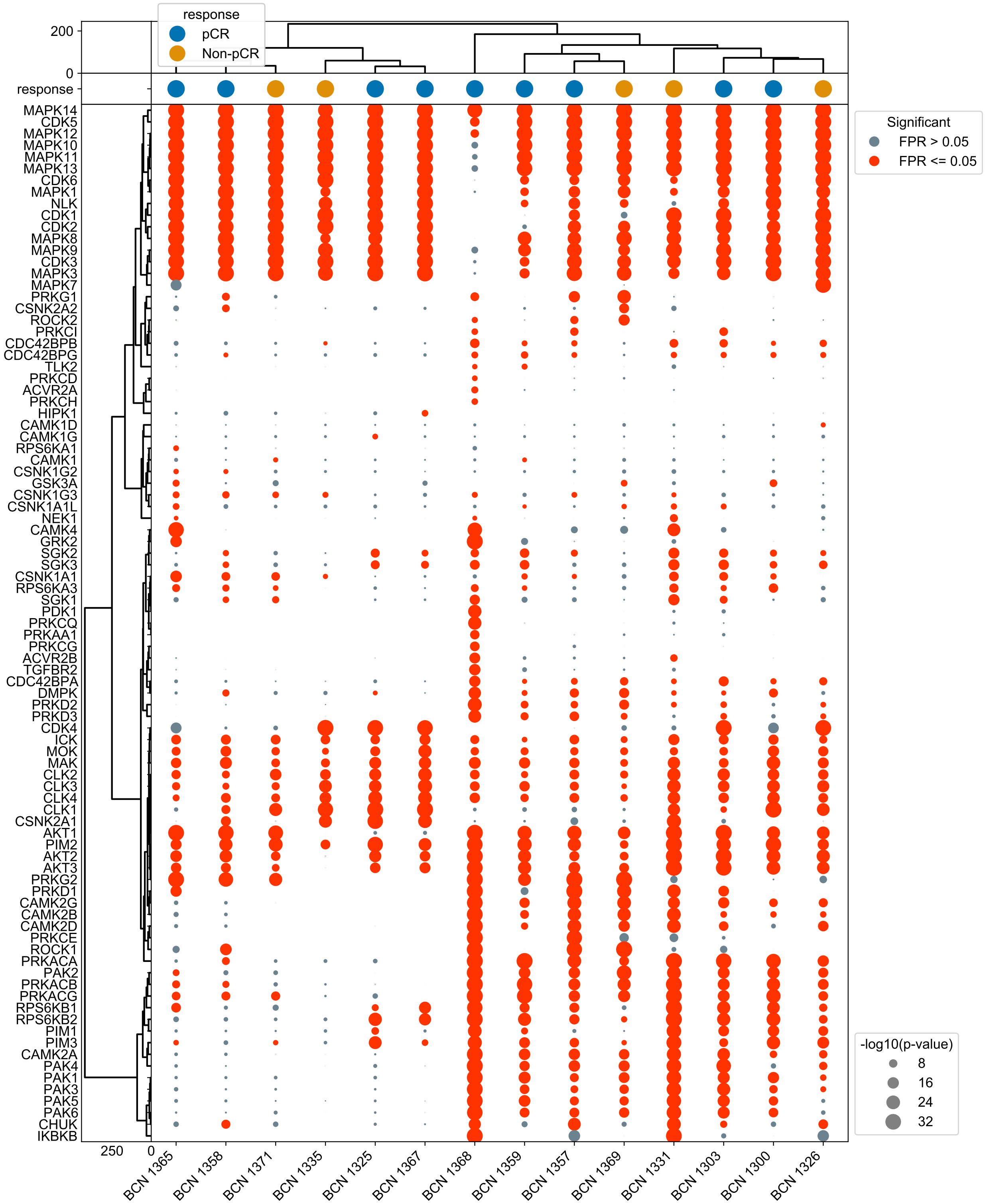


**Figure S6.6. KSTAR serine/threonine kinase activity predictions on microscaled biopsies prior to treatment** Full dotplot containing all serine/threonine kinase activity predictions (for which at least one sample patient had on microscaled biopsies of HER2+ patients for only pre-treatment. Corresponds to the samples from Figure S6.5. Dot size corresponds to the activity score (-log10(p)), and dots are colored based on if the observed activity score has a false positive rate below 0.05. Kinases and samples were sorted using hierarchical clustering with ward linkage. Treatment response to combination therapy of chemotherapy and anti-HER2 therapy (responder/pCR vs. non-responder/non-pCR) of each sample are indicated with the context dots above the dotplot.

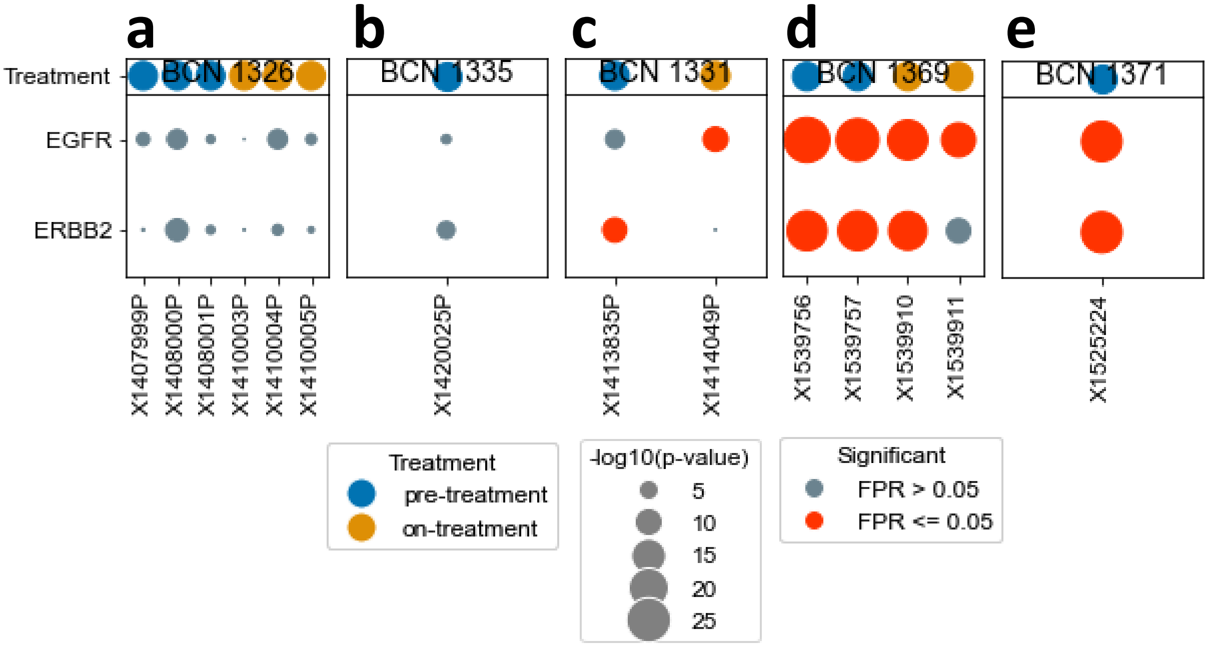


**Figure S6.7. Patient-specific KSTAR predictions for non-pathologically complete responders (non-pCR)** Patient specific dotplots of EGFR and ERBB2 KSTAR activity predictions for non-pathologically complete responders (non-pCR) to treatment with chemotherapy and anti-HER2 therapy. Each plot includes all available replicates and both pre/post-treatment samples. A) Patient BCN1326, a false positive without ERBB2 copy number amplification. B) Patient BCN1331, a pseudo-false positive with ERBB2 amplification but without increased protein expression. C) Patient BCN1335, a pseudo-false positive with ERBB2 amplification but without increased protein expression. Post-treatment data was not available. D) Patient BCN1369, who failed to respond to treatment despite having ERBB2 amplification and increased protein expression. E) Patient BCN1371, who failed to respond to treatment despite having ERBB2 amplification and increased protein expression. Post treatment data was not available.

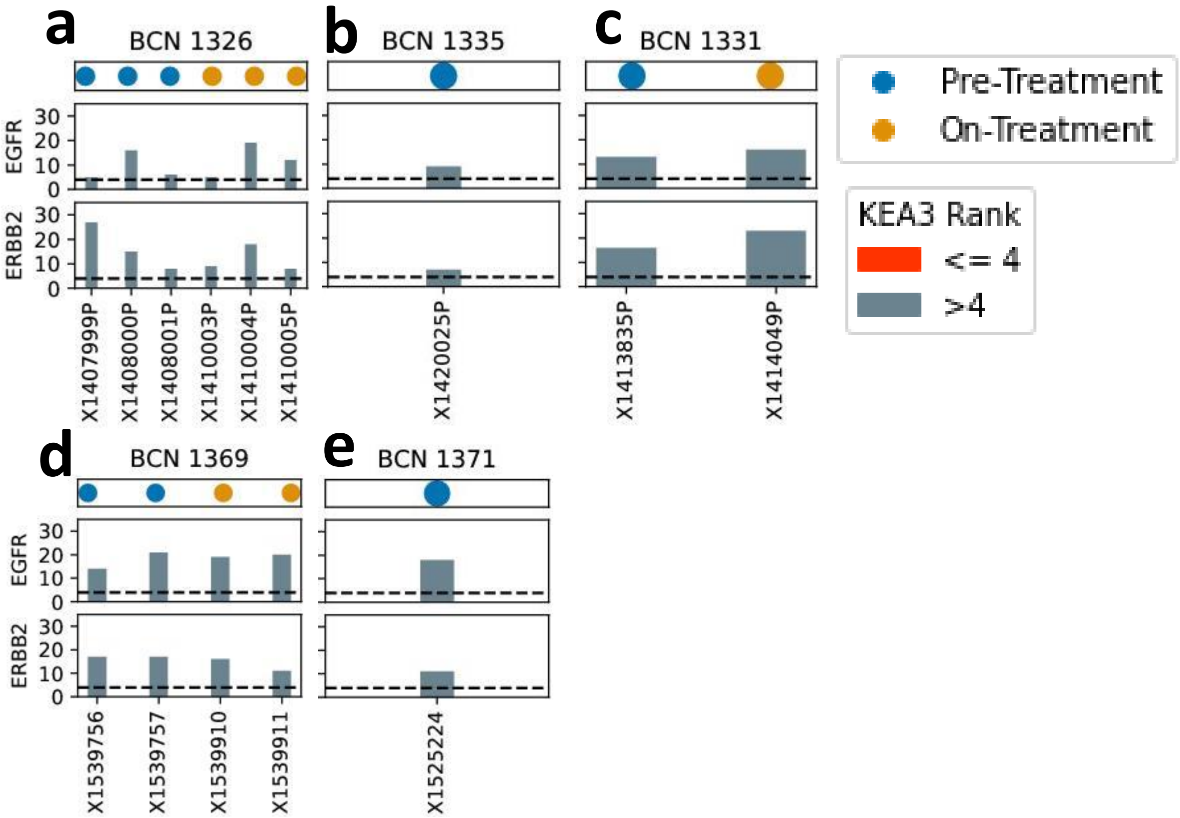


**Figure S6.8. Patient-specific KEA3 ranks for non-pathologically complete responders (non-pCR)** Patient specific dotplots of EGFR and ERBB2 KEA3 ranks for non-pathologically complete responders to treatment with chemotherapy and anti-HER2 therapy. Ranks are relative to the 50 tyrosine kinases found in NetworKIN in order to make ranks comparable to KSTAR. Each plot includes all available replicates and both pre/post-treatment samples. As all but one true HER2+ sample (not identified as a false positive by Satpathy et al (Nature Communications, 2020)) had ERBB2 as one of the top 4 most active kinases based on KSTAR predictions, we have applied a significance cutoff of rank 4 (rank is 4 or better, kinase is deemed significantly active). See Figure S6.11 for more details on this cutoff choice. A) Patient BCN1326, a false positive without ERBB2 copy number amplification. B) Patient BCN1331, a pseudo-false positive with ERBB2 amplification but without increased protein expression. C) Patient BCN1335, a pseudo-false positive with ERBB2 amplification but without increased protein expression. Post-treatment data was not available. D) Patient BCN1369, who failed to respond to treatment despite having ERBB2 amplification and increased protein expression. E) Patient BCN1371, who failed to respond to treatment despite having ERBB2 amplification and increased protein expression. Post treatment data was not available.

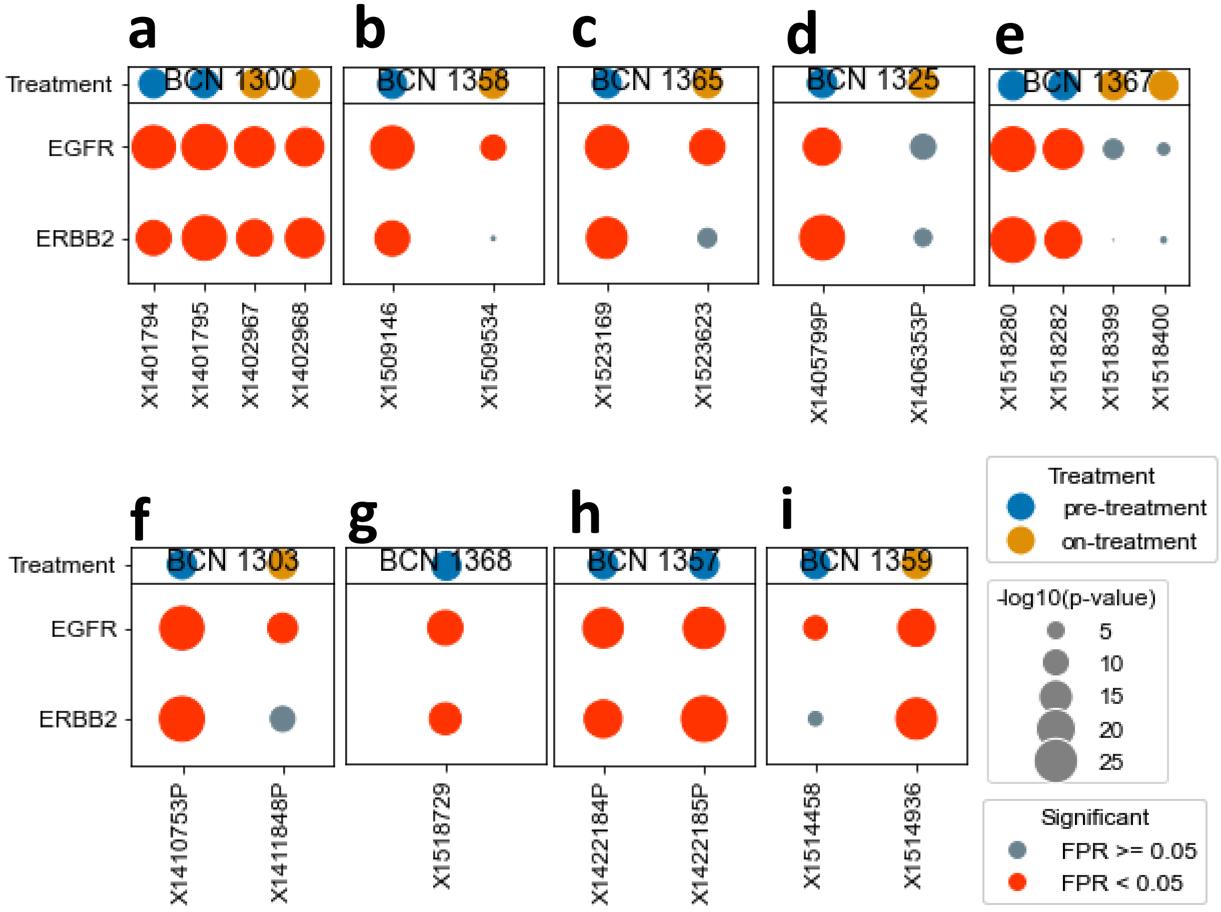


**Figure S6.9. Patient-specific KSTAR predictions for pathologically complete responders (pCR)** Patient specific dotplots of EGFR and ERBB2 KSTAR activity predictions for pathologically complete responders (pCR) to treatment with chemotherapy and anti-HER2 therapy. Each plot includes all available replicates and both pre/post-treatment samples. A) Patient BCN1300. This is the only pCR patient who had basal ERBB2 activity but did not see activity decrease post-treatment, indicating that ERBB2 therapy arm was not responsible for successful response. B) Patient BCN1358. C) Patient BCN1365. D) Patient BCN1325. E) Patient BCN1367. F) Patient BCN1303. G) Patient BCN 1368. Post treatment data not available. H) Patient BCN1357. Post treatment data not available. I) Patient BCN1359. This patient exhibited an interesting response to treatment, where ERBB2 mRNA levels actually increased upon treatment, perhaps explaining the increase in predicted activity.

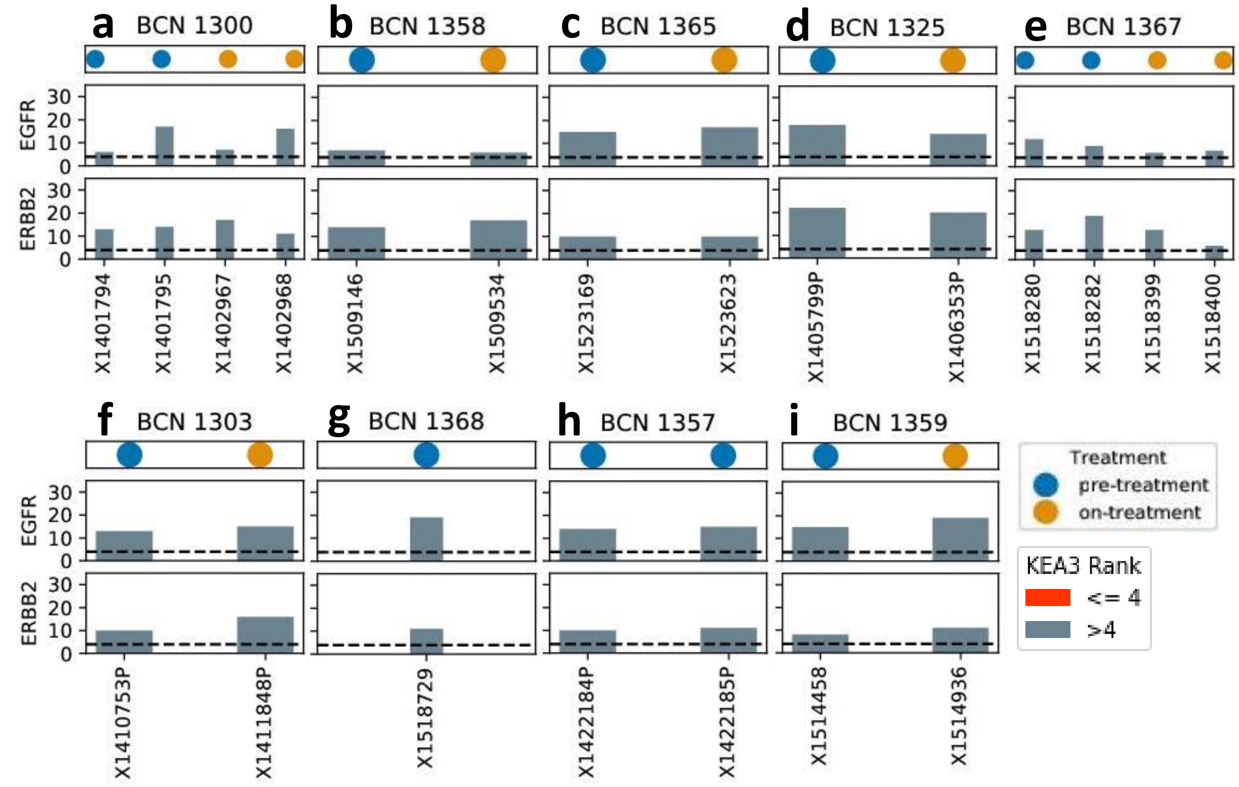


**Figure S6.10. Patient-specific KEA3 ranks for pathologically complete responders (pCR)** Patient specific dotplots of EGFR and ERBB2 KEA3 ranks for non-pathologically complete responders to treatment with chemotherapy and anti-HER2 therapy. Ranks are relative to the 50 tyrosine kinases found in NetworKIN in order to make ranks comparable to KSTAR. Each plot includes all available replicates and both pre/post-treatment samples. As all but one true HER2+ sample (not identified as a false positive by Satpathy et al (Nature Communications, 2020)) had ERBB2 as one of the top 4 most active kinases based on KSTAR predictions, we have applied a significance cutoff of rank 4 (rank is 4 or better, kinase is deemed significantly active). See Figure S6.11 for more details on this cutoff choice. A) Patient BCN1300. This is the only pCR patient who had basal ERBB2 activity but did not see activity decrease post-treatment, indicating that ERBB2 therapy arm was not responsible for successful response. B) Patient BCN1358. C) Patient BCN1365. D) Patient BCN1325. E) Patient BCN1367. F) Patient BCN1303. G) Patient BCN 1368. Post treatment data not available. H) Patient BCN1357. Post treatment data not available. I) Patient BCN1359. This patient exhibited an interesting response to treatment, where ERBB2 mRNA levels actually increased upon treatment, perhaps explaining the increase in predicted activity.

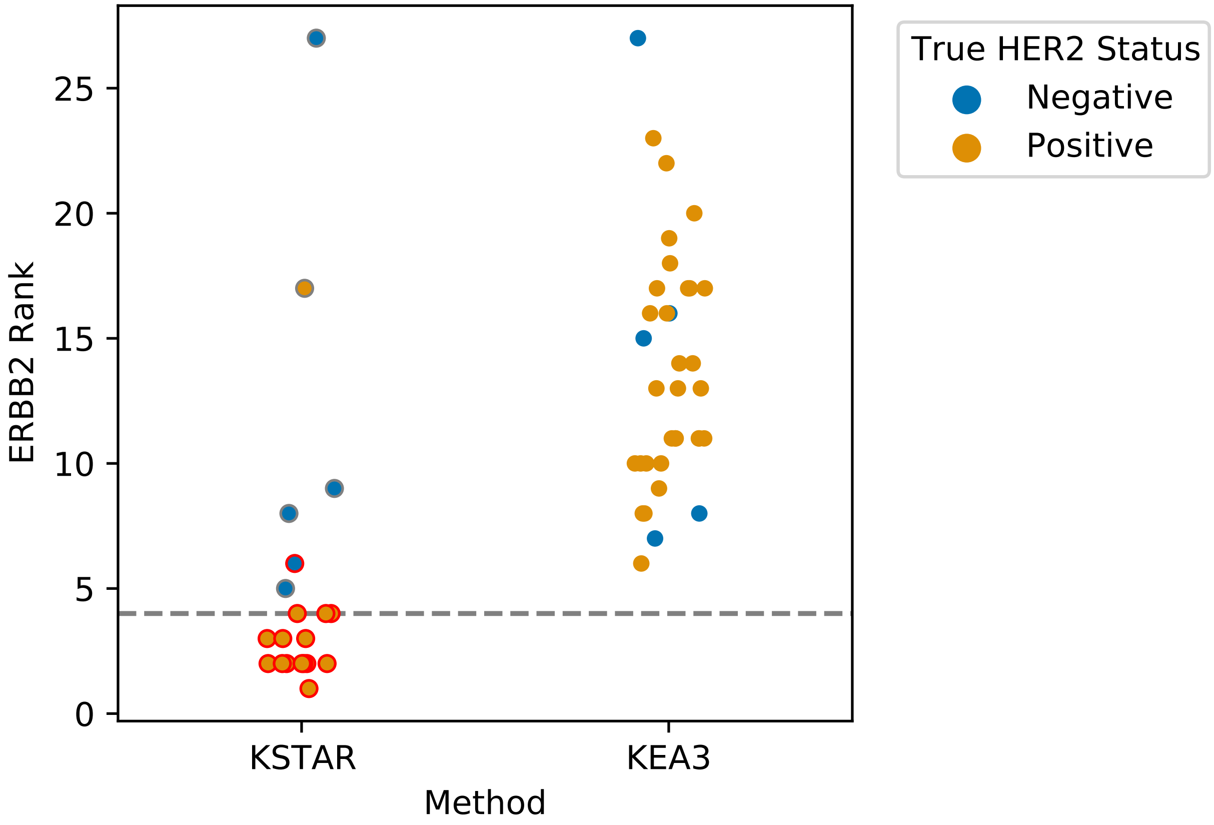


**Figure S6.11. Comparing HER2/ERBB2 activity ranks in KSTAR and KEA3 for microscaled biopsies** The activity rank of HER2/ERBB2 in the pre-treatment microscaled biopsies of HER2+ patients from Satpathy et al. (Nature Communications, 2020), obtained using either KSTAR activity scores or KEA3 mean ranks. Patients are colored according to their true HER2 positivity status identified in Satpathy et al. (BCN1326, BCN1331, BCN1335 were all identified as false positives, and therefore are colored as negative in this plot). All patient replicates are included. As KSTAR uses false discovery rate as the indicator of activity, KSTAR predictions where $FDR <= 0.05$ are indicated with a red edge. Lastly, all but one true HER2 positive patient sample, BCN1359, identified ERBB2 as one of the top 4 most active kinases. Notably, ERBB2 did not appear in the top 4 most enriched kinases in any KEA3 predictions, and there does not appear to be a clear distinction between the ranks of the true postives and the negative samples.
